# No passive sidekick: mitochondrial compensatory evolution during adaptation of ETC mutant lines in *Caenorhabditis elegans*

**DOI:** 10.64898/2026.09.05.749583

**Authors:** Vaishali Katju, Jean-Loup Claret, Zachary P. Dietz, Suzanne Estes, Ulfar Bergthorsson

## Abstract

Proper mitochondrial function is reliant on favourable mitonuclear epistatic interactions between two genomes, the mitochondrial (mtDNA) and the nuclear with contrasting rates of mutation, copy-number and modes of replication and transmission. The compactness and limited coding capacity of animal mtDNA along with their high mutation rates, uniparental inheritance and lack of recombination has contributed to the view that mtDNA is prone to accumulating deleterious mutations, with mitonuclear adaptation primarily attributed to the nuclear genome. Employing a prospective evolution framework in *Caenorhabditis elegans*, we investigated the adaptive capacity of the mtDNA in ETC-deficient lines bearing four focal ETC mutants across three different breeding systems following 60 generations of adaptation. The mtDNA genome can adapt rapidly to mitochondrial dysfunction caused by deleterious mutations in nuclear genes. Nonsynonymous changes in mtDNA protein-coding genes had significantly higher frequencies and were positively correlated with fitness, indicative of positive selection. Mitochondrial gene *nd-1* emerged as a key driver of compensatory mitonuclear adaptation. Notably, compensatory mutations reach higher frequency when they affected ETC subunits in the same complex as the original deleterious mutation. Our results argue for a disproportionately outsized role for mitochondria in mitonuclear adaptation despite their diminutive genome size and protein-coding capacity.

## Introduction

The mitochondrial (mtDNA) and nuclear (nucDNA) genomes of eukaryotes comprise a bi-genomic system whose coordinated functioning is vital not only to metabolism but a slew of other essential cellular processes including cell death, immunity, inflammation, stress responses and signalling (Shen et al. 2022) with implications for aging and disease (Mottis et al. 2019). Pervasive epistatic interactions contribute to this longstanding mitonuclear interdependency, perturbations of which are expected to compromise electron transport chain (ETC) and other critical functions with the potential for deleterious fitness consequences (Rand et al. 2004). As such, this bi-genomic system is particularly amenable for investigating the origin and maintenance of epistatic interactions and the molecular and functional bases of mitonuclear coadaptation. Maintaining genetic compatibility between the eukaryotic mtDNA and nuclear genomes can be challenging for several reasons. First, the mtDNA and nuclear genomes are replicated independently of one another and have different modes of inheritance, both of which can contribute to genetic conflict (Cosmides and Tooby 1981). Second, the spontaneous mutation rates of metazoan mitochondrial genomes can be orders of magnitude higher than their nuclear counterparts (Haag-Liautard et al. 2008; Konrad et al. 2017). Third, the presence of multiple copies of mitochondria within a cell impinges on their population biology with selection and genetic drift capable of operating both at the intra- and inter-individual levels (Rand 2001). Fourth, modern metazoan mitochondrial genomes are characteristically attenuated with respect to size and protein-coding capacity. The radically different genomic architecture, biology and mode of inheritance of these two genomes can lead to divergent evolutionary dynamics and trajectories that may amplify mismatches between the faster-evolving mtDNA and slower-evolving nuclear genomes. How do these two genomes coevolve to maintain their sophisticated crosstalk and a coordinated genetic network necessary for maintaining homeostatic balance despite their divergent genome architecture and evolutionary pressures?

A change in the environment or a fixation of an allele that is detrimental to optimal mitochondrial function sets the stage for mitonuclear compensatory coevolution. It has been argued that a high spontaneous mutation rate and lack of recombination (reduced effective population size) renders mtDNA vulnerable to the accumulation of deleterious mutations via Muller’s ratchet (Lynch et al. 1993). However, there is no evidence for a functional decline in animal mitochondria over evolutionary time (Radzvilavicius et al. 2017). This paradox is addressed by the nuclear compensation hypothesis (Rand et al. 2004; Havird and Sloan 2016) which argues that, given the limited protein-coding capacity of mitochondrial genomes, the nuclear genome is the source of novel compensatory mutations that ameliorate the accumulation of deleterious mutations in the mtDNA genome. Whereas each of the canonical 13 protein-coding genes of a typical metazoan mtDNA genome encode ETC components, the mitochondrial proteome conservatively comprises upwards of 1,000 proteins involved in other essential biochemical functions (Roger et al. 2017; Rath et al. 2021). As such, the nuclear genome is predicted to serve as the primary source of mitonuclear coadaptation.

Several comparative studies have sought to test the hypothesis of an overarching role of the nuclear genome in compensating for mitochondrial dysfunction. These typically involve investigating the rates of molecular evolution of nuclear genes comprising the mitochondrial proteome that participate in oxidative phosphorylation (OXPHOS) (Bogenhagen et al. 2018) by either (i) directly interacting with mitochondrial gene products, or (ii) functioning as subunits of the mitoribosome, or (iii) nuclear-encoded aminoacyl tRNA synthases that participate in translation by loading amino acids on tRNAs in the mitochondrion. Although the results from the comparative approach are often consistent with the nuclear compensation hypothesis, they have not been conclusive (Zhang and Broughton 2013; Sloan et al. 2014; Havird and Sloan 2016; Li et al. 2017; Hill 2020). For example, a correlation in rates of evolution between mitochondrial- and nuclear-encoded mitochondrial proteins can result from their shared relaxation of selective constraints (Zhang and Broughton 2013) rather than an increased rate of fixation of nuclear mutations compensating for an increased rate of deleterious mitochondrial mutations.

A multilevel selection model has been used to explain the lack of erosion of mitochondrial function over time with purifying germline selection on mitochondrial variants in addition to inter-individual selection (Stewart et al. 2008a, 2008b; Camus and Dhawanjewar 2023). Germline selection in combination with multiple copies of mtDNA and higher mutation rate could, in principle, also facilitate faster adaptation in mtDNA relative to nuclear genes for mitochondrial function (Rand 2008). It can be a challenge to differentiate between cause and effect in associations that have emerged over long evolutionary periods using a retrospective evolutionary analysis approach. Multiple outstanding questions regarding the molecular signature of mitonuclear coevolution and adaptation beg further attention. Can the mtDNA genome capacitate rapid mitonuclear adaptation despite its vestigial stature in most metazoans? Specifically, do mitochondrial-encoded genes have a direct role in the arena of compensatory molecular evolution during mitonuclear adaptation? Is there an outsized role of certain gene(s); i.e., are there any major molecular players? Do certain classes of mutations play a predominant role in mitonuclear adaptation, especially compensatory evolution? What are the structural and functional effects of compensatory or adaptive mitonuclear mutations? How does the mating system impinge on the pace of mitonuclear adaptation, if at all?

A prospective experimental evolution approach is a valuable complementation to classical comparative analyses of coevolutionary processes in mitonuclear adaptation. Furthermore, the model nematode *Caenorhabditis elegans* has emerged at the forefront of mitonuclear research given its amenability to experimental evolution coupled with large-scale genome sequencing of replicate experimental populations, *in vivo* approaches to investigate mitochondrial physiology and morphology, ease of conducting RNAi-mediated gene knockdown, treatment with mitotoxicants and the genetic ability to manipulate breeding systems, among others (Estes et al. 2023). We conducted a laboratory evolution experiment to investigate the molecular and functional bases of mitonuclear adaptation in *C. elegans*. *C. elegans* strains bearing four deleterious ETC mutations (two mitochondrial-encoded, two nuclear-encoded) were subjected to laboratory adaptation for 60 generations in large replicate populations under three breeding systems (obligate selfing, facultative outcrossing or obligate outcrossing). The ETC genes comprised the mitochondrial *cox-1* and *ctb-1*, the nuclear *gas-1* and *isp-1* as well as an *isp-1* IV; *ctb-1* M mitonuclear double mutant. Following the adaptation regime, 128 control and experimental populations were subjected to phenotypic analyses for evolution of fitness (Dietz et al. 2025), male frequency and sperm size (Bever et al. 2022). Herein we investigate mtDNA mutations that arose during adaptive recovery that may serve to ameliorate the deleterious effects of the original ETC mutations while assessing the role, if any, that mitochondrial-driven compensatory evolution may play in evolutionary rescue.

## Results

### Identification of 116 mtDNA variants following whole-genome sequencing of adaptation lines subjected to an experimental evolution regime of population expansion

We sought to investigate the genetic basis of mitonuclear adaptation in *C. elegans* by employing an experimental evolution approach that involved subjecting four ETC mutant strains (**Table 1**) to a regime of strong selection via experimental maintenance at large population sizes (*Ne* = 1,000 worms per generation). Moreover, *C. elegans* is unique among experimental metazoan systems given the ability to genetically manipulate sex determination and hence the breeding system, a feature we capitalized upon to further investigate the influence of breeding system on the rate of adaptation. Mutant strains of four electron transport chain (ETC) genes *—* two mtDNA-encoded (*cox-1* and *ctb-1*) and two nucDNA-encoded (*gas-1* and *isp-1) —* under three breeding systems (the wildtype facultatively outcrossing, obligately selfing and obligately outcrossing) (**Supplementary Fig. S1**) were subjected to this adaptative regime for 60 consecutive generations (**Supplementary Fig. S2**), yielding 96 experimental lines (4 ETC mutants × 3 breeding systems = 12 treatments × 8 biological replicates). In addition, we evolved eight replicate lines of a mitonuclear double mutant *isp-1 IV; ctb-1 M* in the wildtype N2 (facultatively outcrossing) breeding background. Lastly, eight replicate lines were evolved as large population size control treatments for each of the three background breeding systems without any associated ETC mutants (total 24 lines). Together, this yielded 128 evolved experimental evolution lines (16 treatments × 8 biological replicates) (see **Supplementary Fig. S1** for experimental design and abbreviated notations for the various treatments and replicate line names). The three breeding systems were designated the notations N, F and X corresponding to a facultatively outcrossing (wildtype N2), obligately outcrossing (*fog-2* deletion mutant) and obligately selfing (*xol-1* deletion mutant), respectively. To further simplify notation, we designate the ETC mutants as *cox*, *ctb*, *gas* and *isp* corresponding to *cox-1*, *ctb-1*, *gas-1* and *isp-1*, respectively. The eight biological replicate lines within each treatment were designated the numbers 1−8. As an example, the eight adaptative recovery lines of the *isp-1* ETC mutant lines within the facultatively outcrossing, obligately outcrossing and obligately selfing breeding system background are referred to as N*isp*.1− N*isp*.8, F*isp*.1− F*isp*.8 and X*isp*.1− X*isp*.8, respectively.

**Table 1.** A description of the affected ETC complex and major phenotypes for each allele (all nonsynonymous mutations) of four focal genes from which congenic progenitor strains were generated.

| ETC-deficient mutant | ETC location | Major phenotype |
| --- | --- | --- |
| *ctb-1* | Complex III: mtDNA-encoded *cyt b* | Delayed reproduction and reduced Complex III activity; reduced Complex I function due to weakened supercomplex I-III-V |
| *cox-1* | Complex IV: mtDNA-encoded core catalytic subunit | Reduced mitochondrial membrane potential and lifespan; increased mitochondrial matrix oxidant burden |
| *gas-1* | Complex I: nDNA encoded 49 kDA iron sulphur protein subunit | Short-lived, reduced fecundity, oxidative stress sensitive |
| *isp-1* | Complex III: nDNA-encoded Rieske iron sulphur protein | Long-lived, slow growth, reduced fecundity, oxidative stress sensitive |
| *isp-1/ctb-1* | Complex III: nDNA-encoded Rieske iron sulphur protein and mtDNA-encoded *cyt b* | *ctb-1(qm189)* partially supresses the negative effects of *isp-1(qm150)* phenotype via beneficial allosteric effects on Complex I; intermediate fitness relative to single mutants |

We used whole genome DNA sequencing (DNA-seq) to map 116 candidate mtDNA variants across 125 of the 128 adaptive recovery (adaptive RC henceforth) lines (**Supplementary File S1**). Three experimental lines (double mutant N*isp*/N*ctb*.5, F*ctb*.7 and F*gas*.2) were excluded from further analysis due to WGS issues relating to low sequence coverage.

### Breeding system does not influence the frequencies and accumulation of mtDNA variants

We first investigated if the breeding system (obligately selfing *vs*. facultatively outcrossing *vs*. obligately outcrossing) impinged on the accumulation of mtDNA variants and their role in the pace of mitonuclear adaptation following the adaptive recovery regime. Upon pooling the 125 sequenced lines by breeding system, we identified 40, 40 and 36 mtDNA variants in the obligately selfing (X), facultatively outcrossing (N) and obligately outcrossing (F) breeding system background, respectively. There was no significant difference in the accumulation and distribution of mtDNA variant frequencies across the three breeding systems overall (Kruskal Wallis rank sum test: *χ^2^* = 2.52, *d.f.* = 2, *p* = 0.28) (**Fig. 1a**). A nonparametric two-way Scheirer-Ray-Hare test similarly found no effect of breeding system (*H* = 1.91, *p* = 0.38, *d.f.* = 2), no significant interaction between breeding system and the background ETC mutation (*H* = 9.04, *p* = 0.34, *d.f.* = 8), but a significant effect of the background ETC mutation (*H* = 25.41, *p* = 4.2 × 10^-5^, *d.f.* = 4) (**Fig. 1b**). These results are in accord with our expectation that the breeding system should not influence the accumulation of mtDNA variants given the clonal nature of mtDNA inheritance. Hence, the variation in mtDNA variant frequencies between treatments is primarily due to the ETC-deficient mutant genetic background and not the breeding system (**Fig. 1b**).

**Fig. 1.**
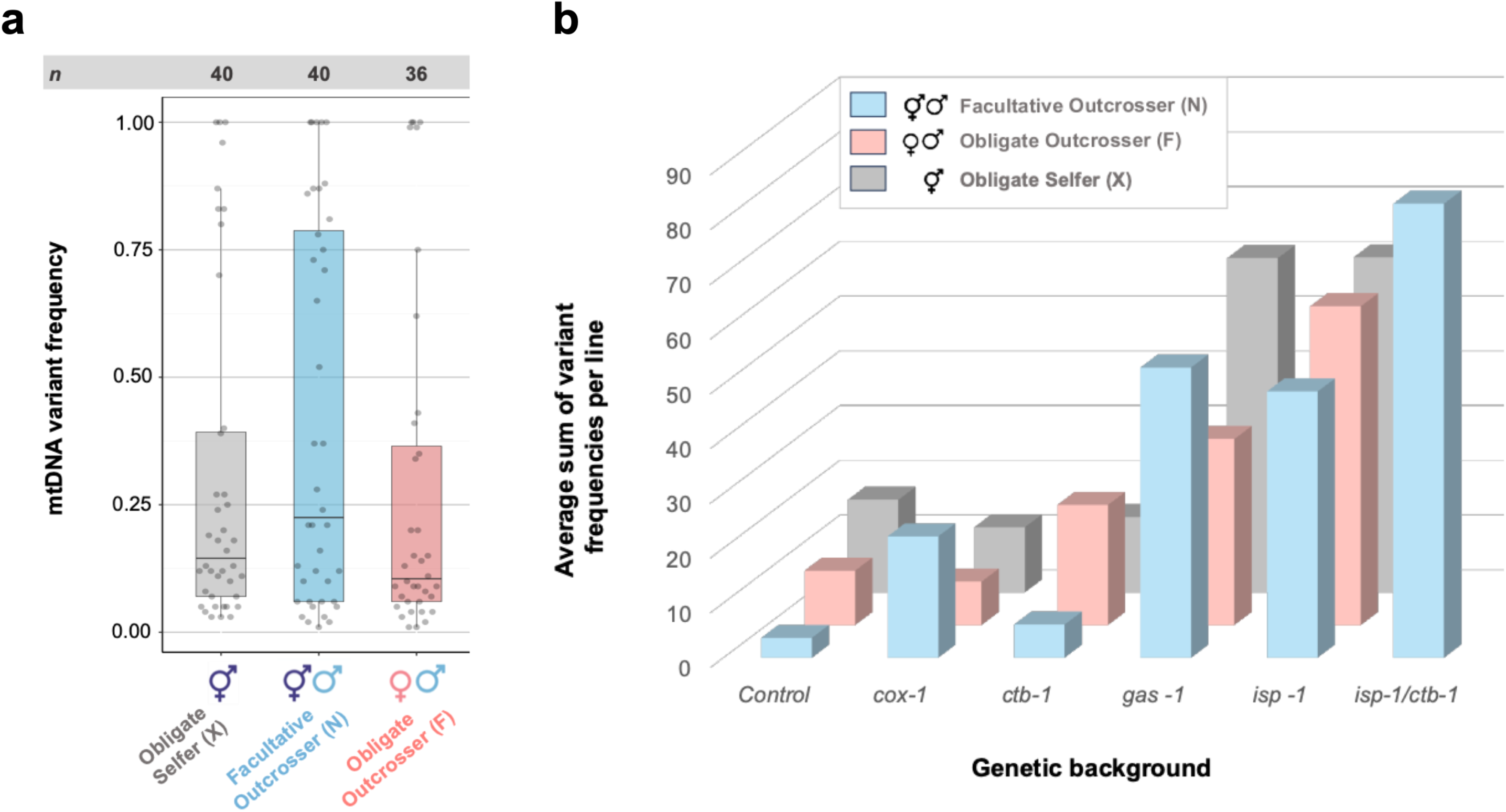
Distribution of mtDNA variants in the adaptive RC lines. **a,** mtDNA variant distribution across the three breeding systems. The pooling of the adaptive RC lines irrespective of ETC mutant or controls yielded 40, 47 and 38 adaptive RC lines within the obligately selfing (X), facultatively outcrossing (N) and obligately outcrossing (F) breeding systems, respectively. The notation “*n*” in the panel on top of the figure denotes the total number of mtDNA variants identified in the experimental lines pooled by the background breeding system. **b**, Average cumulative mtDNA variant frequency per line across the 16 combinations of ETC mutation and breeding system.

### Adaptive RC lines exhibit similar proportions of base substitutions and small indels as those observed in mutation accumulation lines under minimal efficacy of selection

We next investigated if a particular major mutational class was overrepresented in our experimental lines following the adaptive recovery regime by classifying them as either (i) base substitutions, (ii) small indels, (iii) large deletions or (iv) large duplications followed by a comparison to the mutation spectra observed in a long-term *C. elegans* spontaneous mutation accumulation (MA) experiment under minimal selection (Konrad et al. 2017). Base substitutions (also referred to as SNPs), small indels, large deletions and large duplications comprised approximately 39% (45/116), 59% (69/116), 0% (0/116), 2% (2/116) of the newly originated 116 mtDNA variants in our adaptive RC lines. The proportions of SNPs and small indels, two mutational classes that comprise the majority of new variants in both the adaptive RC and MA experiments, were not significantly different (Fisher’s exact test: *p* = 1.00) (**Fig. 2a**). We further tested if the frequencies of synonymous SNPs and indels differed between populations with and without an ETC mutation background. Hypothetically, ETC mutations could influence the mtDNA mutation rate through the differential production of oxygen radicals. There was no discernible difference in the frequency of synonymous mutations and indels between control populations lacking an ETC mutation (*n* = 24) and those bearing an ETC mutation in their genetic background (*n* = 101) (Fisher’s exact test: *p* = 1.00).

**Fig. 2.**
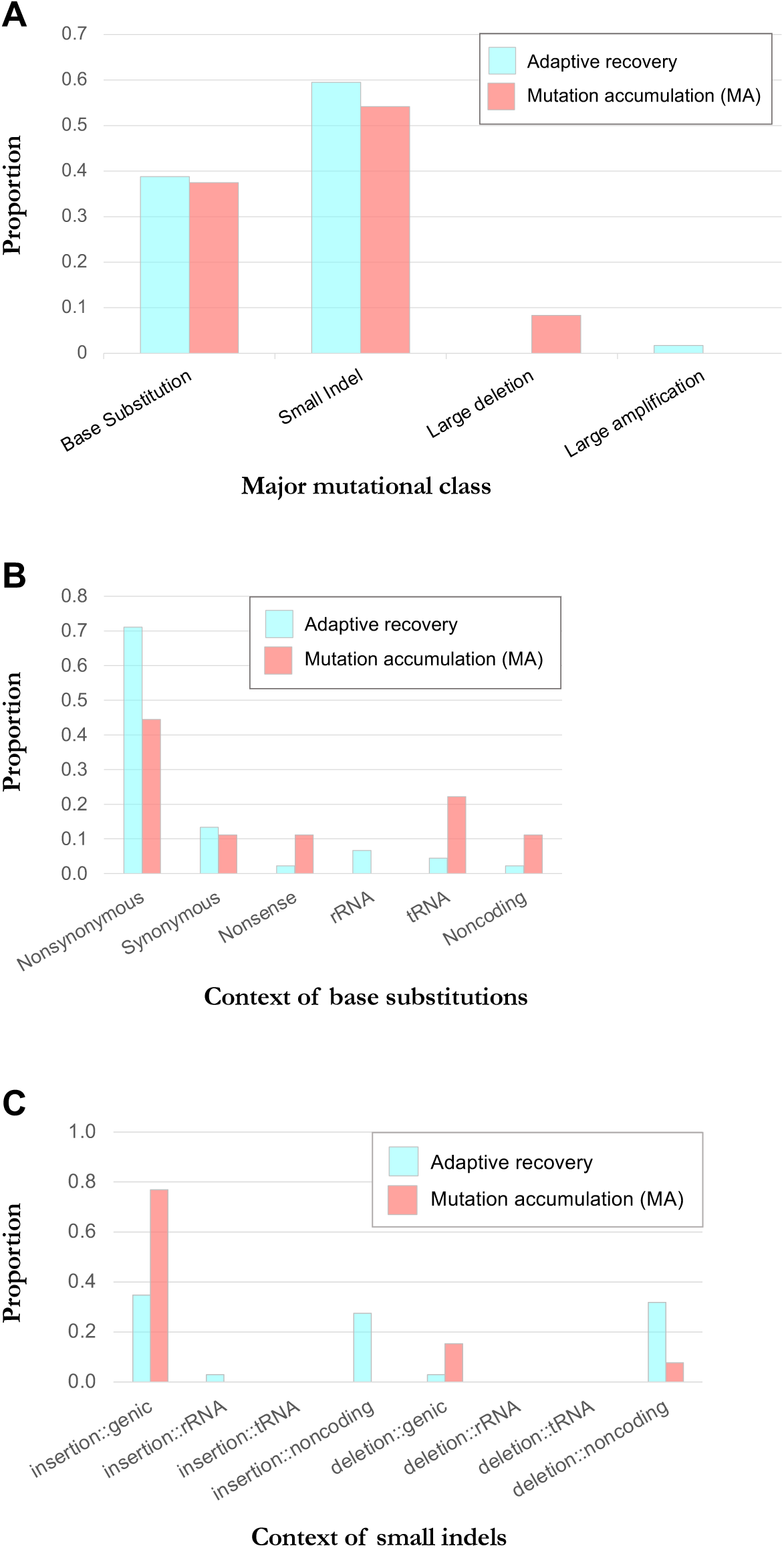
Mutational class and genomic location of new mtDNA variants identified in the adaptive RC lines relative to *C. elegans* MA lines (Konrad et al. 2017). **a**, Proportions of mtDNA variants belonging to four major mutational classes in the adaptive recovery lines in comparison to those observed in MA lines. **b**, Genomic context of base substitutions observed in the adaptive recovery lines relative to MA lines. Base substitutions in protein-coding genes are further classified into nonsynonymous, synonymous and nonsense mutations. **c**, Genomic context of small indels observed in the adaptive recovery lines relative to MA lines. Small indels are further classified into insertions and deletions. “Genic” here refers to mtDNA protein-coding sequences.

The adaptive RC lines lacked large deletions that have been observed in MA experiments and shown to possess selfish properties (Dubie et al. 2020, 2024; Sequeira et al. 2024). Interestingly, an identical large duplication that internally amplified 628 bp within the *nd-5* gene occurred convergently in two adaptive RC lines belonging to different treatments (N*isp*.5 and N*isp*/N*ctb*.1).

### Genomic context of mtDNA base substitutions reveals signature of positive selection in protein-coding genes

Whilst the accrual of small indels and base substitutions in the adaptive RC lines superficially resembles that observed in MA lines, a closer inspection of the genomic context of these two major mutational classes reveals notable divergent characteristics of the adaptation experiment relative to the MA experimental regime. Together, base substitutions comprised 39% of all mtDNA variants in the adaptive RC lines which were further classified based on their genomic location (genic or protein-coding *vs.* rRNA *vs.* tRNA *vs.* noncoding regions) and their proportions compared to their counterparts under MA conditions (Konrad et al. 2017) (**Fig. 2b**). The protein-coding, rRNA, tRNA and noncoding regions correspond to approximately 74%, 12%, 9%, and 5%, respectively of the 13.7 kb *C. elegans* mtDNA genome. The adaptive RC lines had a significantly greater proportion of base substitutions in the protein-coding regions of the mtDNA genome compared to the non-protein-coding regions (rRNA, tRNA and noncoding regions combined) relative to expected based on the proportions of these mtDNA genomic regions (*G*-test statistic = 4.31, *p* = 0.038). The vast majority of base substitutions occurred in protein-coding genes (∼87%), with the predominant class being nonsynonymous changes (∼82%) (**Fig. 2b**). Because nonsynonymous mutations serve as prime targets for engendering beneficial fitness effects during adaptation, their high frequencies are suggestive of a role for positive selection.

### Genomic context of mtDNA small indels reveals signature of purifying selection in protein-coding genes

Small indels represented the largest mutational class in the adaptive RC lines, comprising 59% of all identified new mtDNA variants. This proportion is similar that observed in *C. elegans* MA lines wherein ∼54% of new mtDNA variants were classified as indels (Konrad et al. 2017). As was also previously observed in the MA lines, small insertions exceeded small deletions in the adaptive RC lines by approximately two-fold (65% and 35%, respectively). However, the genomic location of these small indel mutations in the adaptive RC lines is significantly different from their counterparts in the MA lines. Small indels in protein-coding regions can lead to frameshift mutations, resulting in vastly attenuated protein products and/or an altered protein sequence, with deleterious consequences for fitness. We first compared the genomic distribution of small indel mutations in the adaptive RC lines to those observed in the MA lines (**Fig. 2c**). Adaptive RC lines have significantly fewer small indels in protein-coding (genic) regions and significantly more in non-protein-coding regions (tRNA, rRNA and noncoding regions combined) relative to MA lines (Fisher’s exact test: Pearson’s *χ^2^* = 11.72, *p* = 0.0006). In a similar vein, the adaptive RC lines had significantly lower than expected proportions of small indels in protein-coding and RNA genes and greater than expected proportions in noncoding regions (*G*-test statistic = 159.95, *p* = 0.00).

Upon further classification of small indels into insertions and deletion events, it appears that both subclasses of mutations show similar distributions across the different regions of the mtDNA genome. Both insertions and deletions have a significantly sparser distribution in protein-coding genes and significantly greater accumulation in noncoding regions of the adaptive RC lines relative to MA lines (Fisher’s exact test for insertions: *p* = 0.00075; Fisher’s exact test for deletions: *p* = 0.0485). Likewise, the adaptive RC lines had significantly lower than expected proportions of both small insertions and deletions in protein-coding and RNA genes and greater than expected proportions in noncoding regions (insertions: *G*-test statistic = 59.14, *p* = 0.00 46; deletions: *G*-test statistic = 46.71, *p* = 0.00). The observed paucity of small indel mutations in protein-coding and RNA genic sequences suggests the influence of strong purifying (negative) selection.

### Distribution of new variant frequencies in protein-coding genes further exhibits signatures of selection

In addition to the genomic location of the new mtDNA variants, we further analyzed their frequencies within the adaptive RC lines bearing an ETC mutation (*n* = 101). Restricting the analysis to protein-coding genic sequences which comprise ∼74% of the mtDNA genome, we compared the frequency distributions of nonsynonymous and synonymous base substitutions and small indels (**Fig. 3**). The frequency distributions of these three variant classes in protein-coding genes were significantly different (Kruskal Wallis rank sum test: *χ^2^* = 10.80, *p* = 0.0045). The median frequency of nonsynonymous (*n* = 31) and synonymous substitutions (*n* =5) was 75% and 19%, respectively. With the exception of one 3-bp insertion that reached a heteroplasmic frequency of 98.99%, all small indels (*n* = 21) comprised insertions and deletions of 1 or 2 bp, resulting in frameshifts of the ORF and with a median frequency of 12%. Within protein-coding genes, nonsynonymous base substitutions had a significantly higher frequency than small indels (Dunn test with Bonferroni adjustment: test statistic = -3.28, *p* = 0.00314). There was no significant difference between the frequencies of synonymous substitutions and small indels within the protein-coding genes (Dunn test with Bonferroni adjustment: test statistic = -0.90, *p* = 1).

**Fig. 3.**
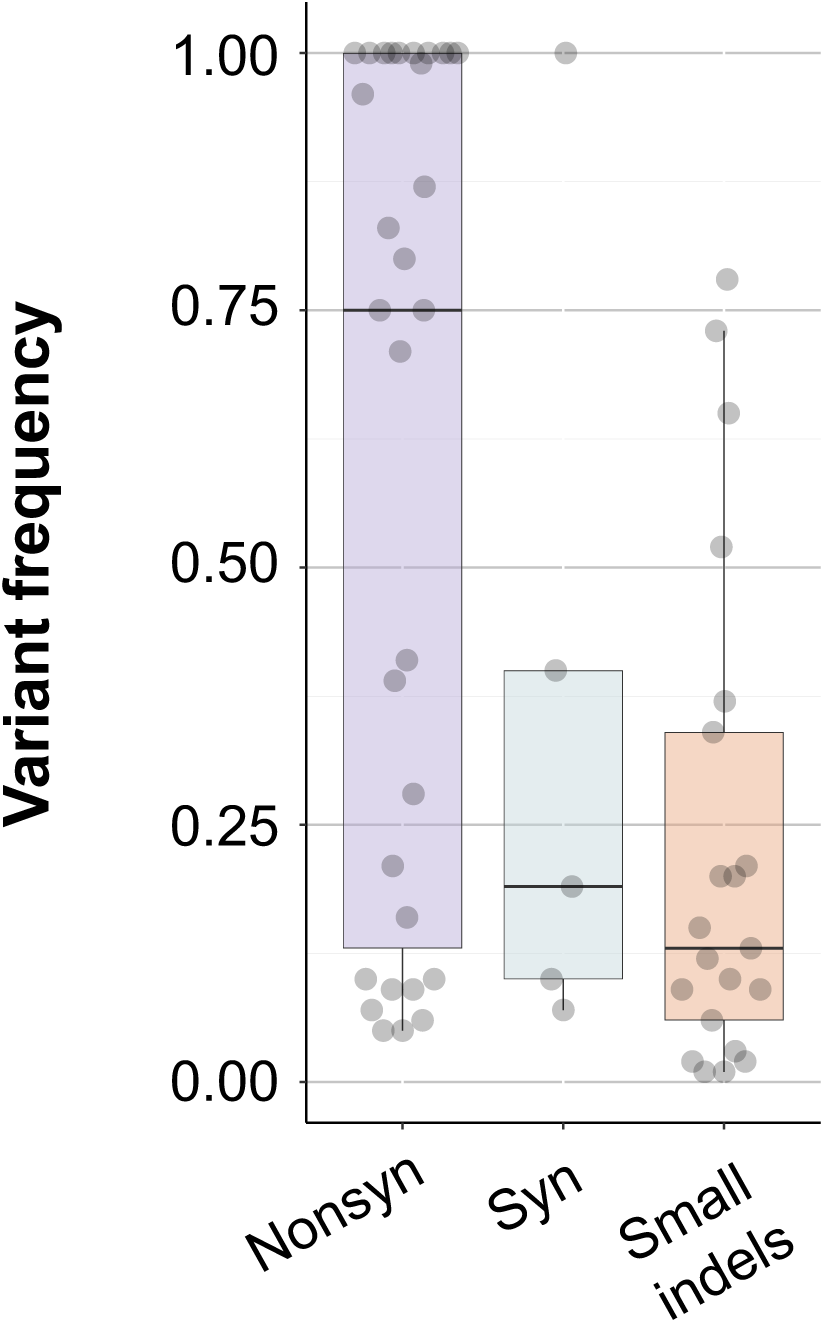
Frequency distributions of two major classes of mtDNA variants (base substitutions/SNPs and small indels) in the protein-coding sequences of the adaptive recovery lines. Base substitutions are further classified into nonsynonymous (nonsyn) versus synonymous (syn) changes. Median frequency is denoted by the horizontal line within the box.

### Distributions of new variants in the adaptive RC lines relative to a mutation accumulation experiment and natural isolates

We additionally investigated the global distribution of mtDNA variant frequencies in our adaptive RC lines relative to their counterparts in a long-term *C. elegans* MA experiment and a previously studied sample of 38 natural isolates (Konrad et al. 2017). First, we pooled all mtDNA variants irrespective of mutational class and compared their frequency distributions. The mtDNA variant frequency distributions of the MA lines, natural isolates and adaptive RC were found to be significantly different (Kruskal Wallis rank sum test: *χ^2^* = 236.73, *p* < 2.2e^-16^) (**Fig. 4a**). The distribution of mtDNA variant frequencies in the MA lines (median: 0.30) and the natural isolates (median: 1.00) represent two distinct spectra of variant frequencies. The mtDNA variants in the MA lines have a uniform frequency distribution spanning low to intermediate to high frequencies whereas the vast majority of mtDNA variants observed in the natural isolates have reached high heteroplasmic frequencies or fixation (homoplasmic) with a smaller subset of extremely low frequency variants. The adaptive RC lines exhibit a distinctly bimodal variant frequency distribution (median: 0.14) with a greater resemblance to the natural isolates than the MA lines with the new variants clustering either in the high- or low-frequency classes.

**Fig. 4.**
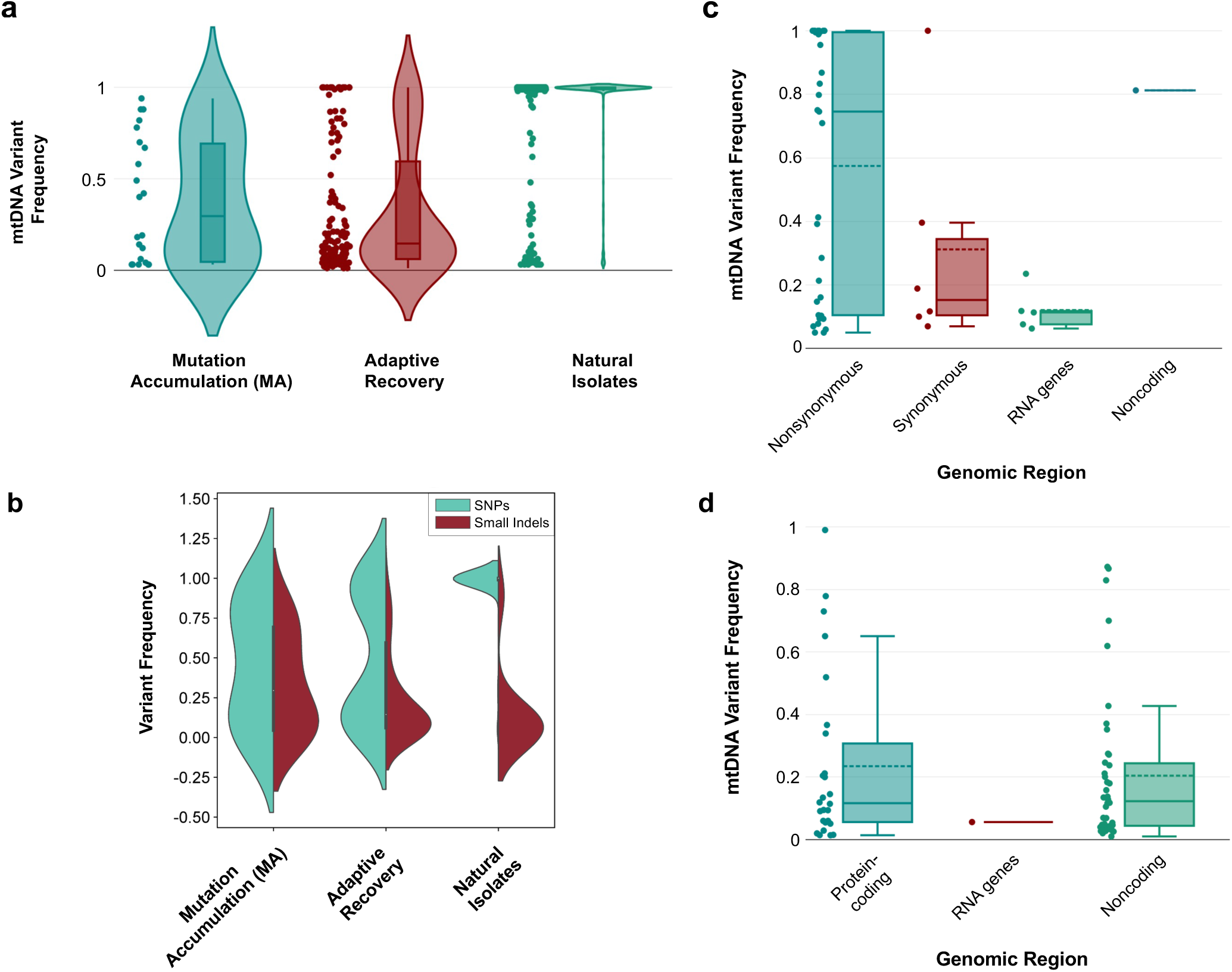
mtDNA variant frequencies in the adaptive RC lines. **a**, Frequency distribution of all mtDNA variants identified in the adaptive RC lines relative to a long-term *C. elegans* MA experiment (Konrad et al. 2017) and 38 natural isolates (Konrad et al. 2017). **b**, Frequency distribution of the two largest mutational classes (SNPs and small indels) of mtDNA variants identified in the adaptive RC lines relative to a long-term *C. elegans* MA experiment (Konrad et al. 2017) and 38 natural isolates (Konrad et al. 2017). **c**, Frequency distribution of SNPs in the adaptive RC lines by genomic location. RNA genes include two rRNA and 22 tRNA genes. Base substitutions in protein-coding genes are further classified into nonsynonymous and synonymous changes. Horizontal solid and dashed lines within boxes represent the median and mean variant frequency, respectively. **d**, Frequency distribution of small indels in the adaptive RC lines by genomic location. RNA genes include two rRNA and 22 tRNA genes. Horizontal solid and dashed lines within boxes represent the median and mean variant frequency, respectively.

Further distinguishing between base substitutions and small indels, the two major classes of observed mtDNA variants within the adaptive RC lines, reveals key differences in their evolutionary dynamics (**Fig. 4b**). In the natural isolates, which are expected to be under the influence of stringent natural selection, the majority of base substitutions and small indels are present in high and low frequency, respectively. Base substitutions may serve more readily as targets for adaptive evolution via positive selection. In contrast, small indels, with their propensity to induce frameshift mutations, are likely to be more deleterious and subject to purifying selection and their presence in low frequencies likely reflects heteroplasmic variation due to the constant input of new mutations. In contrast, MA lines exhibit approximately uniform frequency distributions for both base substitutions and small indels, given that most mutations accumulate neutrally due to the regime of minimal selection efficacy that is the hallmark of such experiments. The adaptive RC lines show a unimodal distribution of small indels at low frequencies, a pattern similar to that observed in the natural isolates. While random mutations provide a constant source of new small indels variants in the mtDNA genome as heteroplasmies, purifying selection can counteract the deleterious phenotypic consequences of such mutations by removing or reducing their frequencies such that they remain below the mitochondrial threshold effect. In contrast to the pattern observed for small indels, the natural isolates exhibit a unimodal distribution of base substitutions in high frequencies (**Fig. 4b**). The adaptive RC lines display a bimodal distribution of SNP variants at both high and low frequencies, with a paucity of new variants at intermediate frequencies. This distribution is suggestive of positive selection driving beneficial variants to higher frequencies while low-frequency SNP variants comprising a pool of (i) newly arisen neutral or beneficial SNP variants at low heteroplasmic frequencies that have not had time to rise appreciably in frequency by drift or selection and (ii) deleterious mutations that are kept at low heteroplasmic frequencies by individual or intra-individual selection.

We further analyzed the frequency distribution of all SNP and small indel variants in our adaptive RC lines in relation to their mtDNA genomic location (**Figs. 4c** and **4d**). With respect to the pool of 45 identified mtDNA base substitutions, 39, five and one variant(s) were observed in protein-coding, RNA and noncoding regions, respectively. Of the 39 base substitutions in the protein-coding sequences, 32, six and 1 variant(s) were nonsynonymous, synonymous and nonsense substitutions, respectively (**Fig. 4c**). Nonsynonymous mutations, aside from being the largest class (71%) of observed base substitutions, had also attained the highest frequencies (median 0.75; range 0.05-1.0) relative to synonymous mutations (median 0.15; range 0.07-0.40) and base substitutions in RNA genes (median 0.11; range 0.06-0.24).

Of the 69 observed small indels, 26, one and 42 variant(s) were observed in protein-coding, RNA and noncoding regions, respectively (**Fig. 4d**). Given that RNA genes comprise 21% of the mtDNA genome, the paucity of small indel mutations (*n* = 1) in RNA genes, and with a low heteroplasmic frequency of 0.056 (5.6%), is especially striking. The vast majority of small indels (61%) are restricted to noncoding regions which only comprises 5% of the mtDNA genome, and with a low median frequency of 0.12 (range 0.01-0.87). Whereas 26 small indels (38%) are observed in protein-coding regions which comprise 74% of the mtDNA genome, the median frequency of these mutations is similarly low (0.12) with a range of 0.01-0.99.

### Fitness recovery of adaptive RC lines is positively correlated with the number of nonsynonymous base substitutions

Our preceding analyses demonstrated that the vast majority of base substitutions (87%) occurred in protein-coding genes, and of these, the vast majority were nonsynonymous mutations (82%). Furthermore, the median heteroplasmic frequency of nonsynonymous mutations was 0.75 and far exceeded that for synonymous mutations (0.19). Both of these features suggest positive selection for nonsynonymous mutations during the adaptive period. Indeed, the fitness increase in the adaptive RC lines (Dietz et al. 2025) was positively correlated with the sum of their heteroplasmic frequencies of nonsynonymous mutations (Pearson correlation coefficient *r* = 0.59, *p* < 0.00001) (**Fig. 5**). In contrast, there was no correlation between the accrual of all other mutations and fitness change during the adaptive regime (Pearson correlation coefficient *r* = 0.01, *p* = 0.91) (**Fig. 5**).

**Fig. 5.**
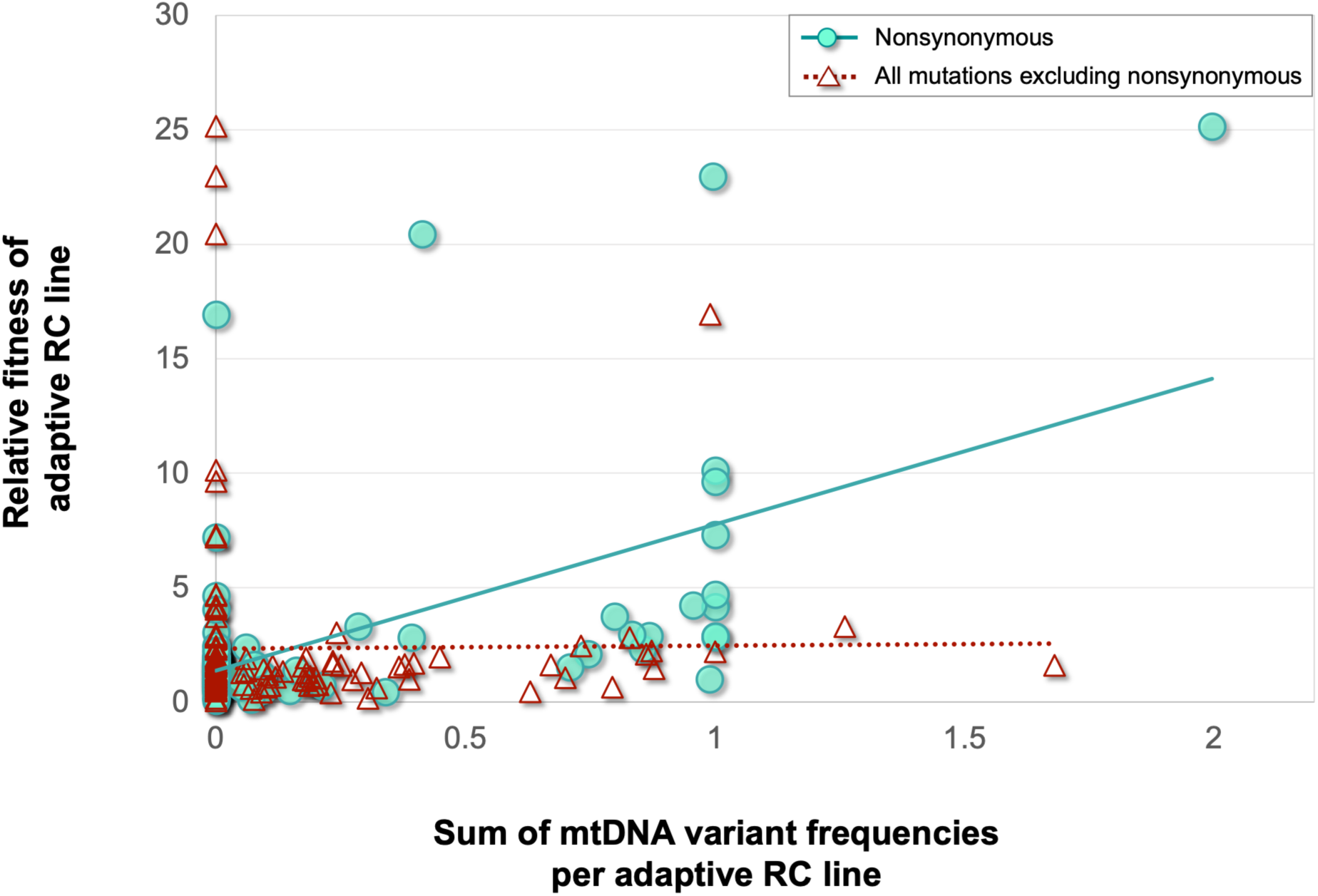
Change (evolved/ancestral) in relative fitness as a function of cumulative variant frequency per line. Productivity of adaptive RC lines following 60 generations of population expansion was used as a proxy for fitness (Dietz et al. 2025). There is a significant positive association between fitness increase and nonsynonymous variant frequency (filled turquoise circles) but not to other mtDNA variant frequencies (open maroon triangles).

### High frequency nonsynonymous mutations occur in mtDNA genes belonging to the same ETC complex as the background ETC-deficient mutation

As the frequency of nonsynonymous mutations is associated with an increase in fitness, we further asked if nonsynonymous mutations that occur in ETC mutant lines had the same frequencies when they occurred in an ETC mutant of the same or different respiratory complex. The original ETC mutants affect three different respiratory complexes, ETC complex I (*gas-1*), III (*isp-1*, *ctb-1*) and IV (*cox-1*). The spontaneous nonsynonymous mutations that arose during 60 generations of evolution occurred in mitochondrial genes coding for proteins in ETC complex I (*nd-1*, *nd-2*, *nd-4*, *nd-5* and *nd-6*), III (*ctb-1*), IV (*cox-1*, *cox-2*) and V (*atp-6*). The within-line frequencies of spontaneous nonsynonymous mutations were significantly higher in mtDNA genes localizing to the same ETC complex as the background ETC-deficient mutation than their counterparts in genes belonging to a different ETC complex from the background ETC-deficient mutation (Mann-Whitney *U* = 23, *z* = -3.57, *p* = 0.00036) (**Fig. 6a**). The average and median frequency of spontaneous nonsynonymous mutations that arose in mtDNA genes belonging to the same ETC complex as the background ETC-deficient mutation was 0.96 and 1.00, respectively. In contrast, the average and median frequencies of such mutations occurring in mtDNA genes belonging to a different ETC complex as the background ETC-deficient mutation was 0.36 and 0.10, respectively. Eight *gas-1* (ETC complex I) mutant lines had high-frequency nonsynonymous mutations (0.75-1.00) in the genes *nd-1* and *nd-6*, both mtDNA-encoded subunits of ETC complex I (**Fig. 6b**). Similarly, three of these mutations in the *isp-1* mutant background (ETC complex III) occurred in *ctb-1*, which also localizes to ETC complex III (**Fig. 6c**). Each of these three nonsynonymous *ctb-1* mutations reached fixation during the adaptation experiment. One of these *ctb-1* mutations is identical to a *ctb-1* mutation that has been previously described as partially compensating for the nuclear *isp-1* mutation (Feng et al. 2001). The far greater population frequency of mutations when they occur in the same ETC complex as the original ETC mutation suggests that compensatory mutations often involve direct physical interaction of proteins−but not necessarily directly interacting amino acid residues.

**Fig. 6.**
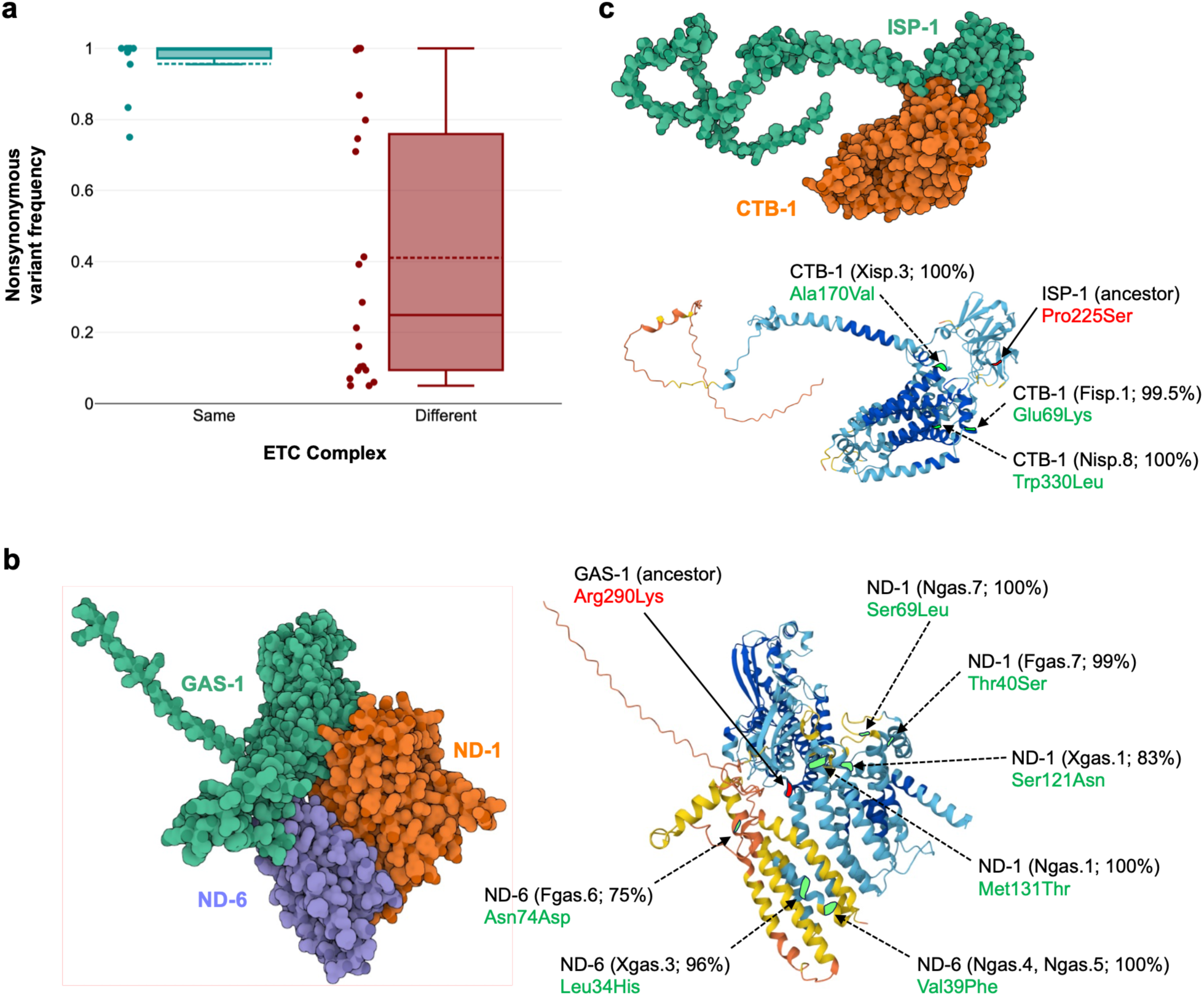
Frequencies and locations of nonsynonymous mtDNA mutations in the adaptive RC lines. **a,** Nonsynonymous mtDNA mutations reached higher frequencies when they arose in genes whose protein products localize to the same ETC complex as the original ETC-deficient background mutation (teal) compared to those localizing to a different ETC complex (salmon). Horizontal solid and dashed lines within boxes represent the median and mean variant frequency, respectively. **b,** Predicted interactive structure of the nuclear GAS-1 protein and mtDNA proteins ND-1 and ND-6 in ETC complex I accompanied with the predicted locations of the ancestral deleterious ETC-deficient *gas-1*(*fc21*) mutation and putative compensatory high-frequency mutations in ND-1 and ND-6. The illustrative left panel depicts the predicted structural interaction between the GAS-1, ND-1 and ND-6 proteins in ETC Complex I. The right panel displays the quaternary structure with the arrows pointing to the predicted location of the ancestral ETC-deficient *gas-1*(*fc21*) mutation (red) and the putative compensatory mtDNA nonsynonymous mutations (green) with the name of the mtDNA gene bearing the new variant, the adaptive RC line(s) and the frequency of the new variant in parentheses, and the amino acid residue position and direction of amino acid replacement. **c,** Predicted interactive structure of the nuclear ISP-1 protein and mtDNA protein CTB-1 in ETC complex III accompanied with the predicted locations of the ancestral deleterious ETC-deficient *isp-1*(*qm150*) mutation and putative compensatory high-frequency mutations in CTB-1. The illustrative left panel depicts the predicted structural interaction between the ISP-1 and CTB-1in ETC Complex III. The right panel displays the quaternary structure with the arrows pointing to the predicted location of the ancestral ETC-deficient *isp-1*(*qm150*) mutation (red) and the putative compensatory mtDNA nonsynonymous mutations (green) with the name of the mtDNA gene bearing the new variant, the adaptive RC line(s) and the frequency of the new variant in parentheses, and the amino acid residue position and direction of amino acid replacement.

### Disproportionately higher incidence of nonsynonymous mutations in nd-1

Given that the vast majority of base substitutions (87%) occur in protein-coding genes and comprise nonsynonymous mutations (82%), we next sought to investigate the genic distribution of these mutations. Do we observe an outsized role of a certain mtDNA gene or a subset of mtDNA genes during molecular adaptation? Rather than an even distribution of mutations across the 12 protein-coding mtDNA genes, a striking pattern was emergent. Of the 32 nonsynonymous mutations that arose in the adaptive RC populations, 14 (44%) were located in *nd-1* (**Fig. 7**) despite *nd-1* comprising only 9% of the protein-coding sequence of the *C. elegans* mitochondrial genome (*G* = 28.36, *p* < 10^-6^).

**Fig. 7.**
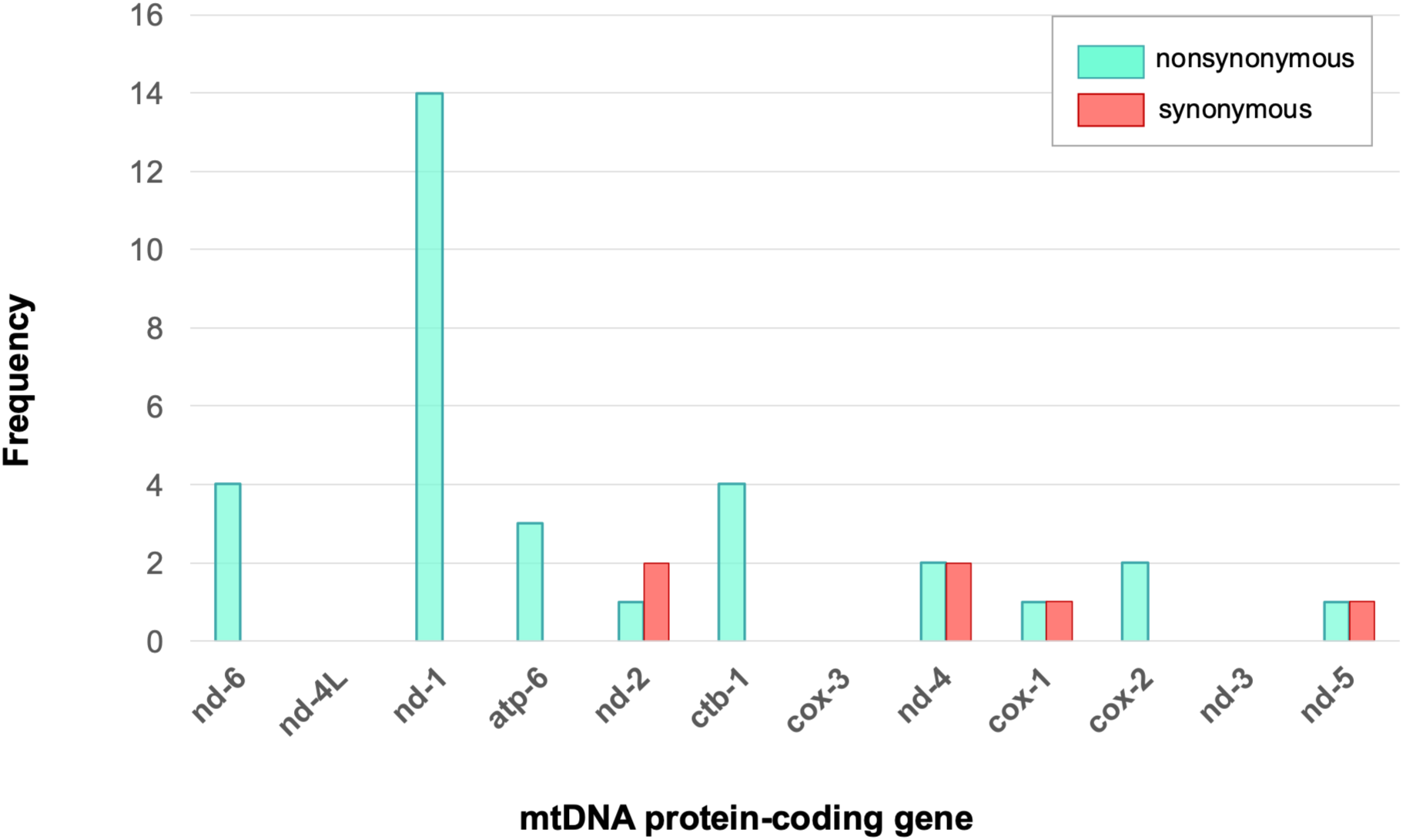
Base substitutions per protein-coding gene. Frequency distribution of novel base substitutions in the adaptive RC lines across the 12 protein-coding genes comprising the *C. elegans* mtDNA genome. Base substitutions are further classified into nonsynonymous (red) and synonymous changes (green).

Although there is significant overrepresentation of high frequency or fixed mutations in mtDNA genes localizing to the same ETC complex as the gene bearing the background ETC-deficient mutation, an interesting exception is the case of the adaptive RC lines bearing the background *isp-1* ETC-deficient mutation. The majority of the *nd-1* mutations (ETC complex I) arose in *isp-1* background mutants (ETC complex III). Adaptive RC lines with the original *isp-1* deficient mutation incurred 19 base substitutions, 18 of which were nonsynonymous. However, the vast majority of nonsynonymous mutations (13 of 18) in these *isp-1* (ETC complex III) ETC-deficient mutant lines occurred in mtDNA-encoded subunits of ETC complex I (genes *nd-1*, *nd-2* and *nd-4*). The mtDNA gene *nd-1* was overrepresented as a target for nonsynonymous base substitutions in the *isp-1* mutant lines, harbouring 10 of the 18 mutations (*G* = 24.93, *p* < 10^-6^) with an average heteroplasmic frequency of 42%. In addition, *nd-1* also incurred an in-frame 3 bp insertion with an heteroplasmic frequency of 99% but no synonymous mutations. The disproportionately high number of nonsynonymous mutations in *nd-1* suggests selection for variants compensating for the original *isp-1* mutation. Two additional nonsynonymous mutations in genes for ETC complex I in the *isp-1* lines occurred in *nd-4* where they reached frequencies of 99% and 79% in 60 generations.

### nd-5 is a hotspot for indels

We plotted the genic distribution of 27 insertion/deletion events including the two large internal gene amplifications (**Supplementary Fig. S3**). Small genic insertions (23 variants) vastly exceeded small indels (two variants). Hence as was observed under an MA regime with minimal selection (Konrad et al. 2017), the *C. elegans* mtDNA genome has an insertion bias. A particularly surprising observation was the high incidence of frameshift mutations in *nd-5*. All but one of these *nd-5* frameshift mutations were either a +1 or +2 bp insertion in a homopolymeric run of eight Ts towards the 5′ end of the gene at mtDNA position 11,721. A single +1 bp frameshift was similarly observed in a second homopolymeric run of eight Ts just downstream at position 11,777. Both of these sites were previously found to be hotspots for frameshift mutations in an MA experiment (**Supplementary Fig. S4**; Konrad et al. 2017). The +1 bp frameshift in *nd-5* at location 11,721 is predicted to result in a severely truncated 18 aa peptide whereas the +2 bp insertion results in a 15 aa peptide. The *nd-5* frameshift mutations had an average frequency of 8% in three of the 24 adaptive RC lines without an ETC mutation in their genetic background but an average frequency of 20%, 17%, and 66% in four of 24 *cox-1*, four of 23 *ctb-1*, and five of seven N*isp-1/*N*ctb-1* ETC-deficient mutant lines, respectively. Except for the double mutant N*isp-1/*N*ctb-1* lines, most of the *nd-5* frameshift mutations did not exceed 20% frequency within lines. In contrast, the high frequency distribution of *nd-5* frameshifts in the *isp-1/ctb-1* lines was similar to what has been observed during a long-term MA experiment (**Supplementary Fig. S4**; Konrad et al. 2017). No *nd-5* frameshift mutations were detected in the 23 lines with a *gas-1* ETC-deficient mutation.

In addition to the simple frameshift mutations, *nd-5* also incurred internal duplications of 628 bp at the same location in two lines, N*isp/*N*ctb.1* and N*isp.5*. The breakpoints of the duplication occur in an 8-bp repeat (TATTTTTT) followed by the introduction of a premature stop codon just downstream of the first breakpoint, and predicted to result in a 368 aa peptide sequence compared to the full-length wildtype protein sequence of 527 aa. Both of these duplications reached high frequency of 95% within their respective populations. The history of the strains and the presence of the diagnostic *ctb-1* mutation in one of the populations suggests that, despite the identical locations and near identical frequencies, these duplications arose independently.

### Did mitochondrial dysfunction modify the rate and spectrum of mtDNA mutations?

It has been postulated that high levels of ROS in mitochondria have a major impact on the rate and spectrum of mtDNA mutations and contribute to the relatively high mutation rate in animal mtDNA (Harman 1972; Miquel et al. 1980; Martin and Palumbi 1993). If elevated mitochondrial ROS increases the mtDNA mutation rate, this could have consequences for how rapidly compensatory mtDNA mutations can accrue. Similarly, a modified mtDNA spectrum that increases transversions relative to transitions is predicted to elevate the proportion of nonsynonymous mutations at two-fold degenerate sites (Cheng et al. 1992). Cumulatively, the spectrum of mtDNA mutations in our adaptive recovery *C. elegans* populations was indistinguishable from a large study on the mutation spectrum in experimental *C. elegans* lines under relaxed selection (Fisher’s exact test, two-tailed: *p* = 0.86) (**Supplementary Fig. S5**; Waneka et al. 2021). However, the spectrum of mtDNA mutations in our adaptive RC lines differs significantly from the spectrum of mtDNA polymorphism in the wild (Fisher’s exact test, two-tailed: *p* = 5.87 × 10^-8^) (**Supplementary Fig. S5**; Schifano et al. 2025). The *gas-1* mutation has been shown to increase mitochondrial ROS (Kayser et al. 2001, 2004), which is predicted to increase the frequency of G/C→T/A transversions. The relative number of these transversions in the *gas-1* adaptive RC lines is indeed higher (46%) than in the non-*gas-1* genetic backgrounds (25%), but the differences are not significant (Fisher’s exact test: *p* = 0.29). However, it remains a possibility that the other genetic backgrounds used in this study may have altered the rate and spectrum of mtDNA mutations in a way that contributed to their dynamics of compensatory evolution.

### Identification of potential altORF candidates that overlap with high frequency mutations

Although animal mitochondrial genomes are very compact with limited intergenic noncoding DNA, it has been hypothesized that they also code for alternative open reading frames (altORFs) or small ORFs (smORFs) encoding small functional proteins within genes. For example, an altORF that encodes a 99-amino-acid peptide within the human mitochondrial gene *nd4* appears to contribute to cell and mitochondrial physiology (Kienzle et al. 2023). To consider the possibility that new mtDNA variants arising within our adaptive RC lines may have occurred in such altORFs, we conducted the first characterization of their potential locations within the *C. elegans* mtDNA genome. We identified 229 putative mtDNA altORFs (including smORFs) (**Supplementary Fig. S6a, Supplementary File S2**)—96 forward and 133 reverse—using an *in silico* approach. Among these, the OpenProt (v. 2.2) database returned 15 forward altORF candidates that overlapped the following mitochondrial genes: known coding regions as follows: l-rRNA (3), s-rRNA (1), *cox-1* (2), *nd-4* (2), overlapping *nd-1* (2), *nd-4L* (2), *ctb-1* (2) and *atp-6* (1) (**Supplementary Fig. S6b**). Four of these hypothetical altORFs overlapped 10 mtDNA variants at nine sites in the adaptive RC lines (**Supplementary File S3**). Eight of these 10 variants were found in *nd-1*, and one each in the *atp-6* and *ctb-1* genes (**Supplementary Fig. S6b**). All of these mutations were characterized as either nonsynonymous substitutions or containing an in-frame single amino acid insertion. More work would be required to determine whether any of the hypothetical altORFs generate functional products and, if so, whether any variants they acquired contributed to adaptive evolution in our RC lines. Because our study suggests a substantial capacity for variants within canonical mtDNA genes to drive adaptive evolution, there may be little reason to invoke a major role for altORFs in such evolution.

## Discussion

Mitochondrial function is a fundamental evolutionary compromise between two genomes (mtDNA and nucDNA) with contrasting rates of evolution and modes of replication and transmission. This compromise is reliant on the coordinated expression of thousands of nuclear genes and a handful of mtDNA genes still residing in the vestigial mitochondrial genome. Because most ETC components are subject to this dual genetic control, the maintenance of favourable mitonuclear epistatic interactions is key to proper mitochondrial functioning (Rand et al. 2004; Osada and Akashi 2012). At the cellular level, mitonuclear coordination is mediated by a sophisticated bi-directional communication system comprising anterograde and retrograde signals. From an evolutionary perspective, we are interested in how this genomic coevolution is maintained given the radically different population biologies and baseline mutational rates experienced by nucDNA and mtDNA. It can be a challenge to differentiate between cause and effect in associations that have emerged over long evolutionary periods using a retrospective evolutionary analysis approach. Multiple outstanding questions regarding the molecular signature of mitonuclear coevolution and adaptation beg further attention. We employed a prospective evolution framework in *C. elegans* to investigate the adaptive capacity of the vestigial mitochondrial genome in ETC-deficient lines experiencing mitochondrial dysfunction.

### Evidence for positive selection in mtDNA protein-coding genes with a disproportionately outsized role of nonsynonymous mutations

We first address the question as to whether the animal mtDNA genome with its limited size and protein-coding capacity has a role, if any, in capacitating mitonuclear adaptation. Specifically, do mitochondrial-encoded genes have a direct role in the arena of compensatory molecular evolution during mitonuclear adaptation? Additionally, does a certain class of mtDNA mutations play a predominant role in compensatory mitonuclear adaptation as evidenced by fitness recovery of lines bearing deleterious ETC-deficient mutations? Many features of mitochondrial biology would appear to promote high rates of deleterious mutation accumulation compared to the nuclear genome including maternal inheritance, minimal or no recombination (reduced effective population size, *Ne*), and limited DNA repair. Indeed, a preeminent model on the subject, the *nuclear compensatory hypothesis*, is based on the premise that the higher mutation rates of animal mtDNA lead to the accrual of a nonadaptive and deleterious mutation load, which in turn enhances the selective pressure on the nuclear genome to counteract by accumulating adaptive compensatory mutations (Dowling 2014). In other words, the *nuclear compensatory hypothesis* posits that mitonuclear matching is maintained to a large extent by the nucDNA genome. Our experimental design aimed to study the adaptive response to both mitochondrial- and nuclear-encoded deleterious mutations (two each), with the nuclear-encoded deleterious mutations in *gas-1* and *isp-1* engendering greater deleterious fitness effects (Dietz et al. 2025). We found no evidence of any mtDNA mutations during adaptation that qualified as reverse (back) mutations or intragenic mutations in the *ctb-1* and *cox-1* ETC mutant lines following adaptive recovery.

The spectrum of spontaneous mtDNA mutations under minimal efficacy of selection have been comprehensively studied in *C. elegans* via MA experiments with SNPs and small indels comprising the vast majority of novel variants (Konrad et al. 2017; Waneka et al.

2021). The spectrum of the two predominant mutational classes (SNPs and small indels) in the adaptive RC lines is indistinguishable from those observed under a regime of relaxed selection in MA experiments. As such, the supply of novel mutations available for future molecular adaptation in the adaptive RC lines is similar to that observed under MA conditions. Because of the unique biology of mitochondria which exist as populations comprising hundreds of thousands copies within individuals, genetic variation is initially manifested as heteroplasmies with the new variants occurring in extremely low frequencies. Heteroplasmic frequencies can be altered dynamically both intraindividually and intracellularly in response to the evolutionary forces of genetic drift and the operating mode of selection. Purifying selection acts to reduce heteroplasmic frequencies of deleterious mtDNA mutations whereas positive selection can drive up heteroplasmic frequencies of beneficial mtDNA mutations culminating in their fixation (homoplasy). Moreover, mitochondrial quality-control mechanisms can, in principle, act as means for intraindividual selection on novel mtDNA variants, both negative (purifying) and positive. A central result of this study is the clear contrast between the mtDNA mutational input and the selective filtering of that mutational variation during adaptive recovery. The frequencies of these novel mtDNA variants at the culmination of the adaptation experiment reveal a striking signature of selection with the bimodal frequency distribution of these SNPs—clustering at either very high or very low frequencies—contrasting sharply with the uniform distribution seen in MA lines under near neutral conditions. This bimodality is also apparent in natural populations of *C. elegans* that are subject to strong natural selection (Konrad et al. 2017; Schifano et al. 2025).

There exist several initial lines of support for a significant role of mtDNA nonsynonymous base substitutions in the compensatory mitonuclear adaptation of experimental lines, namely: (i) the adaptive RC lines had a significantly greater number of base substitutions in the protein-coding fraction of the mtDNA genome relative to rDNA, tRNA and noncoding regions, (ii) the vast majority (82%) of base substitutions in the protein-coding genes were nonsynonymous changes, and (iii) the nonsynonymous substitutions exhibited a significantly higher median heteroplasmic frequency (75%) relative to the other mutational classes observed in protein-coding regions (synonymous substitutions, 19%; small indels, 12%). Moreover, a major finding of this study is the significant positive correlation between the fitness recovery of lines (Dietz et al. 2025) and variant frequencies of nonsynonymous mutations accrued within the line, but the lack of any such correlation when all other accrued mutations are considered (synonymous mutations in protein-coding regions, other SNP mutations in nongenic regions of the mtDNA genome and small indels across the entire mtDNA genome). This strong positive association between fitness recovery and the frequency of nonsynonymous mutations provides direct evidence that these variants are not merely hitchhiking but are the primary drivers of compensatory adaptation. Together, these results support a major contribution of mtDNA nonsynonymous mutations in mitonuclear adaptation resulting in fitness recovery of ETC-deficient low-fitness lines. Although nuclear DNA likely plays a role in RC line adaptation—and the mtDNA altORF variants we identified may also contribute—it is clear that variants within the canonical mitochondrial genome were the primary drivers. The context-dependent fitness effects of novel mutations are also apparent here. Under standard environmental conditions in the laboratory, nonsynonymous mutations in mtDNA protein-coding appear to have deleterious fitness effects and are subject to strong purifying (negative) selection (Kotrys et al. 2024). However, in an altered biological context involving the genetic background bearing a deleterious mutation and strong selection operating at large population sizes, nonsynonymous mutations are prime targets for positive selection enabling fitness recovery.

Given the randomness of the mutational process and the limited temporal span of our adaptation experiment, it is to be expected that not all of our adaptive RC lines would exhibit fitness gains. Indeed, the adaptive RC could be classified into two broad classes, namely a low-fitness (minimal or no fitness gain) and a high-fitness class (Dietz et al. 2025). Focusing on the latter, 28 adaptive RC lines exhibited a ≥ two-fold increase in fitness relative to their ancestral strain (range 2.09- to 25.15-fold) and of these, 22 (79%) bore at least one mutation in a mtDNA protein-coding gene. Within this subset of 22 adaptive RC lines, 86% (19/22) of the lines had accrued at least one nonsynonymous mtDNA mutation. Notably, the median heteroplasmic frequency of these novel mtDNA nonsynonymous variants was a high 92% with ten having reached fixation (frequency >99%) in a mere 60 generations. Furthermore, adaptive RC line F*isp*.1 exhibiting the greatest fitness gain (∼25-fold) possessed two fixed nonsynonymous mtDNA mutations in *nd-1* and *ctb-1*. There appear to be multiple paths to fitness recovery even within the diminutive 13.8 kb mitochondrial genome of *C. elegans*, as evidenced by the fact that these candidate beneficial mutations appear in different genes, at different sites within genes and sometimes in different ETC complexes from the original deleterious mutation.

### Mitochondrial gene nd-1 emerges as a key driver of compensatory mitonuclear adaptation

A striking observation of this study is the disproportionately outsized role of *nd-1* in steering adaptive evolution of ETC-deficient lines. The 12 mtDNA protein-coding genes of *C. elegans* comprise ∼75% genomic fraction of the total mtDNA genome so mutational input in genic regions is expected. The *nd-1* gene comprises a small fraction of the mtDNA protein-coding genome (∼9%). Yet, it harboured nearly half of all the observed novel nonsynonymous substitutions that arose during the adaptation experiment, particularly in populations evolving from an *isp-1* (ETC complex III) deficient background.

The mitochondrial-encoded protein ND1 (NADH:ubiquinone oxidoreductase core subunit 1) is a critical component of ETC complex I (NADH dehydrogenase) in the mitochondrial electron transport chain. As subunits comprising the first and largest enzyme in the mitochondrial respiratory chain which serves as the major portal for electrons entering the respiratory chain, complex I genes play a central role in energy metabolism. As such, ETC complex I constitutes a rate-limiting step in overall respiration (Sharma et al. 2009). The L-shaped structure of complex I comprises (i) a hydrophilic peripheral arm located in the matrix and composed of nuclear-encoded subunits and (ii) a hydrophobic membrane arm lodged within the inner mitochondrial membrane and composed entirely of mtDNA-encoded subunits (ND1, ND2, ND4, ND4L, ND5 and ND6) (Efremov et al. 2010; Baradaran et al. 2013). The release of redox energy from electron transfer in the peripheral arm of complex I subsequently drives proton pumping in the membrane arm and the coupling of these two steps in complex I are thought to be affected by conformational changes of the two arms (Baradaran et al. 2013; Fiedorczuk and Sazanov 2018). The ND1 protein’s unique location at the base of the peripheral arm and at the junction of the complex’s peripheral and membrane arms places it at the structural and functional "hinge" of the enzyme. ND1 is essential for complex I assembly as it makes direct contact with the nuclear-encoded subunits of the peripheral arm, and houses the quinone-binding chamber while linking it with the proton-pumping modules in the membrane arm (Fiedorczuk and Sazanov 2018). ND1 dysfunction not only impedes the assembly of a mature complex I but also initiates a domino effect in negatively impacting the downstream formation of OXPHOS supercomplexes and the assembly and/or stability of complex IV (Lim et al. 2016).

The ND1 subunit’s critical role in several different functions is perhaps best exemplified in humans wherein a diverse array of diseases/syndromes occur owing to base substitutions in the *nd-1* gene. These mutations are broadly classified into four groups thought to influence (i) quinone accessibility, (ii) coupling of the Q site and proton pumps, (iii) proton-pumping activity of the E channel and (iv) enzyme dysfunction by affecting structural stability (Fiedorczuk and Sazanov 2018). The pivotal position occupied by ND1 at the junction of the peripheral and membrane arms of ETC complex I may enable its action as a primary transducer of conformational changes between these two arms, thereby rendering it an ideal target for compensatory mutations as well as during adaptation. By altering the coupling efficiency at this interface, the cell may be able to recalibrate the entire electron transport chain to offset the downstream bottlenecks caused by mutations perturbing ETC function. The human *nd-1* variant Tyr30His is a case in point, as it has both been associated with Leber hereditary optic neuropathy (Obayashi et al. 1992) as well as high altitude adaptation in Tibetan and Indian populations (Ji et al. 2012). In *C. elegans* natural isolates, *nd-1* is both (i) more polymorphic than other mtDNA protein-coding genes (Schifano et al. 2026) and (ii) a more frequent target for recurrent mutations (Nguyen et al. 2025). One mtDNA site in particular, mtDNA:2024 corresponding to codon 94 in *nd-1* was identified as experiencing episodic diversifying selection (Nguyen et al. 2025).

### Structural proximity and compensatory mtDNA evolution

Further support for compensatory mtDNA evolution in the adaptive RC lines comes from the association between the ETC complex affected by the ancestral ETC-deficient nuclear mutation and the location and frequencies of derived nonsynonymous mtDNA substitutions. Nonsynonymous mutations arising in mtDNA genes encoding subunits of the same ETC complex as the original deleterious nDNA mutation reached higher frequencies than mutations in other complexes, suggesting compensatory fine-tuning within physically and functionally interacting protein assemblies. Of the 10 nonsynonymous mutations observed across nine adaptive RC lines with a *gas-1* deleterious mutant background, eight were in the *nd-1* and *nd-6* genes (median frequency of 0.99) which colocalize with *gas-1* in ETC complex I. Similarly, we observe the repeated fixation of *ctb-1* nonsynonymous mutations in adaptive RC lines with an *isp-1* deleterious mutant background, both colocalizing within ETC complex III. The *isp-1* allele was originally characterized as a mutation that increased life span, albeit with substantially slower developmental rates (Feng et al. 2001). One of the mtDNA mutations that arose in the *isp-1* RC lines (Ala170Val) had previously been characterized as a suppressor of slow development in *C. elegans isp-1* mutants (Feng et al. 2001). This *ctb-1* mutation independently arose and went to fixation in RC line X*isp*.3, underscoring the reproducibility and convergence of these compensatory pathways. In addition, nonsynonymous mutations in *ctb-1* arose and reached fixation in *isp-1* RC lines at two other sites, namely Glu69Lys in F*isp*.1 and Trp329Leu in N*isp*.8. Such patterns are difficult to reconcile with purely indirect or pleiotropic effects and instead point to direct protein–protein or complex-level interactions as key targets of selection. This suggests that the most effective way to restore ETC function is through direct physical interaction or localized structural stabilization within the same complex. Previous studies have found that intragenic compensatory mutations tend to be more effective when they are close to the original deleterious mutation (Poon & Chao 2006; Davis et al. 2009). Our results show that spatial proximity of compensatory mutations to deleterious mutations is also positively associated with their effectiveness in ameliorating the fitness cost of deleterious mutations even for extragenic compensatory mutations.

A notable exception to the above trend is particularly intriguing. There was an abundance of *nd-1* and *nd-4* mutations (complex I) in the *isp-1* deleterious mutant (complex III) background. Of the 14 *nd-1* nonsynonymous mutations observed in our adaptive RC lines, 10 (71%) occurred in *isp-1* RC lines. However, while these nonsynonymous *nd-1* mutations were more numerous than mutations in *ctb-1*, only one of these had reached fixation at the termination of the experiment. The remaining nine nonsynonymous *nd-1* mutations occurred in frequencies ranging from 5−87%. Furthermore, a single amino acid insertion in *nd-1* also reached near fixation (98.9%) in adaptive RC line F*isp*.4. In addition to these *nd-1* mutations, two *isp-1* adaptive RC lines had *nd-4* variants in 80% and 99.5% frequency. The abundance of *nd-1* and *nd-4* (complex I) mutations in the *isp-1* RC lines (complex III) suggest a cross-complex compensatory mechanism, perhaps through the optimization of the broader respiratory supercomplex I·III·IV or a metabolic shift that alleviates the bottleneck at complex III by modulating complex I activity (Acín-Pérez et al. 2008; Suthammarak et al. 2010).

### Functional significance of high-frequency mtDNA nonsynonymous mutations during adaptive evolution

Of the seven high-frequency mtDNA nonsynonymous changes observed in lines with a *gas-1* deleterious mutant background, four mutations and three mutations were observed in *nd-1* and *nd-6*, respectively. Here, both the deleterious mutant and the putative adaptive changes are restricted to complex I. These mutations were compared to *nd-1* and *nd-6* linked diseases in humans (reviewed in Fiedorczuk and Sazanov 2018). Two *nd-1* mutations, Thr40Ser and Ser72Leu, correspond to the *nd-1* region associated with the quinone-binding chamber and therefore quinone accessibility. The other two *nd-1* mutations, Met131Thr and Ser121Asn, correspond to a region involved in creating the proton input channel for the first translocation channel (referred to as the E channel); hence both these mutations are likely involved in the proton-pumping activity of the E channel. The three *nd-6* mutations, Asn74Asp, Val39Phe, and Leu34His, correspond to regions involved in proton translocation, either directly or indirectly.

The nature of high-frequency mtDNA mutations in adaptive RC lines with an *isp-1* deleterious mutant background is more intriguing. As described above, the *gas-1* deficient lines accrued high-frequency nonsynonymous mutations in mtDNA genes (*nd-1* and *nd-6*) belonging to the same ETC complex as the original deleterious *gas-1* mutant (i.e. complex I). In contrast, the *isp-1* deficient lines accrued high-frequency mutations in mtDNA genes belonging to both (i) the same (complex III) and (ii) a different complex (complex I) as the original deleterious *isp-1* mutant (i.e. complex III). Three *ctb-1* nonsynonymous mutations (Glu69Lys, Ala170Val and Trp330Leu) reached fixation in three high-fitness *isp-1* deficient lines. Both the CTB1 and ISP1 subunits are components of complex III. Two *isp-1* adaptive RC lines accrued nonsynonymous mutations in *nd-4* (Ser120Tyr, Ser364Phe). Four *isp-1* adaptive RC lines accrued three nonsynonymous mutations (Pro39Lys, Pro39His, Val215Phe) and one fixed in-frame isoleucine insertion at amino acid residue 154 in the *nd-1* gene.

Interestingly, even though the putative compensatory mutations occur in mtDNA genes of both complex I and III, all three genes contribute to a functional complex I (Fiedorczuk and Sazanov 2018; Suthammarak et al. 2010). While *nd-1* functions pleiotropically, one of its critical functions is to maintain structural stability and hence contribute to the assembly of an intact complex I. Likewise, *nd-4* is implicated in the correct assembly of complex I. And lastly, even though *ctb-1* codes for a complex III subunit, one of the compensatory mutations in this gene appears to ameliorate fitness through improving complex I functions (Suthammarak et al. 2010).

### Molecular signature of intense purifying selection against small indels

In stark contrast to the signature of positive selection on nonsynonymous changes in mtDNA protein-coding regions, small indel variants in protein-coding regions appear to be, on average, under stringent purifying selection. This purifying selection on small indels is manifested in both their (i) biased genomic locations and (ii) frequencies. Whereas base substitutions in the form of nonsynonymous changes are overrepresented in the mtDNA protein-coding regions, small indels (especially frameshift-inducing mutations) are significantly depleted in protein-coding and RNA genes, with the vast majority restricted to noncoding regions. This biased genomic location of small indels in our adaptive RC lines is also in stark contrast to *C. elegans* MA lines which accrued small indels throughout the mtDNA genome (Konrad et al. 2017). Furthermore, even when small indels occurred in protein-coding regions, for example in the *nd-5* mutational hotspot also observed by Konrad et al. (2017) in MA lines, they occurred at low heteroplasmic frequences (median frequency 12% barring one in-frame 3 bp insertion). Nonsynonymous substitutions, on the other hand, attained high heteroplasmic frequencies or fixation. This indicates that while the spontaneous mutation rate for small indels remains high, especially in homopolymeric runs, frameshift-inducing mutations engender a high fitness cost and are therefore prevented from reaching high frequencies, thereby maintaining them as a mutational load below some mitochondrial threshold effect.

We further note the high incidence of small indel mutations at two hypervariable sites in the *C. elegans* mtDNA genome. The first hypervariable site at genomic position mtDNA:3,234 is unusual in that it is a large poly-A sequence, which are very rare on the coding strand of the *C. elegans* mitochondrial genome. The other hypervariable site, a poly-T tract early in the *nd-5* gene at mtDNA:11,721 is clearly a hotspot for indels in both experimental and natural populations of *C. elegans* (Konrad et al. 2017; Schifano et al. 2026). However, there are 28 within-gene poly-T sequences with ≥7 Ts in the *C. elegans* mitochondrial genome, which raises the question of why length variation at this particular site is so common. One possibility is that frameshift mutations at this site contribute to selfish properties through replication/transmission bias of T length variants at this site (Dubie et al. 2024).

Heteroplasmic mtDNA deletions have been found in both laboratory and natural isolates of *C. elegans* and the evidence suggests that they have selfish properties in that their proliferation is associated with lowered fitness (Gitschlag et al. 2016; Dubie et al. 2020, 2024; Sequeira et al. 2024). A few natural isolates of *C. elegans* have mtDNA duplications of the control region and adjacent genes in relatively low heteroplasmic frequency (Schifano et al. 2026). The two high-frequency intragenic duplications in *nd-5* reported here are the first to be described for laboratory populations. The fact that these duplications are in *nd-5* is interesting in light of the fact that many natural populations of *C. briggsae* contain an *nd-5* pseudogene adjacent to their functional *nd-5*, and noncoding regions in *Caenorhabditis* species have resulted from *nd-5* duplications (Li et al. 2018). For example, the mitochondrial genomes of *C. briggsae* and *C. sinica* contain an additional pseudo-*nd-5* located between *nd-2* and *ctb-1* (Li et al. 2018). In an analysis of the acquisition of novel noncoding regions in *Caenorhabditis* mtDNA, *nd-5* was the only genic source of these regions that could be identified (Li et al. 2018). The fact that (i) the internal *nd-5* duplications in lines N*isp*.5 and N*isp*/N*ctb*.2 create a termination codon downstream of the duplication breakpoint, and (ii) these lines do not show significant fitness improvement leaves open the question of whether duplications of certain *nd-5* regions result in a biased transmission of mtDNA containing them, which might also explain why *nd-5* is the only genic source of noncoding regions in *Caenorhabditis* mtDNA.

### Mitochondrial evolutionary dynamics during adaptation are independent of the breeding system

In nature, a transition from outcrossing to a primarily selfing mode of reproduction is sometimes associated with an increase in the accumulation of nonsynonymous mutations which have been presumed to be largely deleterious (Paland and Lynch 2006; Neiman et al. 2010). In contrast, purifying selection is at least as efficient in the mitochondrial genome of some hermaphroditic taxa relative to related sexually reproducing species (Brandt et al. 2017). Our study found no detectable effect of the breeding system—whether obligately selfing, outcrossing, or facultatively outcrossing—on the accumulation or frequency distribution of mtDNA variants. In contrast to the nuclear genome where outcrossing promotes effective recombination and selection efficacy, our observation is consistent with (i) the uniparental transmission of mitochondria and (ii) the clonal nature of the mitochondrial genome with limited or no recombination. Rather, mitochondrial-capacitated adaptation is forged by the internal selective environment of the mitonuclear genotype rather than the organism’s mating system or modulation of the mutation rate.

### Conclusions

The combination of high mutation rates, uniparental inheritance and lack of recombination in animal mitochondrial genomes has contributed to the view that mtDNA is particularly prone the passive accumulation of deleterious mutations and that the adaptive evolution of mitonuclear interactions is primarily attributable to the nuclear genome. The results of experimental evolution in *C. elegans* show that the mitochondrial genome can adapt rapidly to mitochondrial dysfunction caused by deleterious mutations in nuclear genes. The diversity of nonsynonymous mutations that rose to high frequency during compensatory evolution and their association with increased fitness demonstrate a disproportionately outsized role for mitochondria in capacitating rapid mitonuclear adaptation despite their diminutive genome size and protein-coding capacity. Our results suggest that mitonuclear epistasis might be mediated by mtDNA compensatory mutations that are extragenic but co-localized to interacting proteins in multi-protein complexes of the ETC rather than an obvious latch-and-key mechanism between the focal deleterious mutation and putative compensatory mutations. There appears to be an outsized role for mutations in *nd-1* in compensating for both the nuclear *gas-1* and *isp-1* deleterious mutations. Future work will further quantify the contribution of nuclear variants to compensatory evolution in our adaptive RC lines with the potential to identify candidate nuclear genes with hitherto unidentified mitochondrial function. Another exciting avenue awaiting further inquiry is the relative contribution of intra-versus inter-individual selection in dictating the fixation probabilities of compensatory or beneficial mtDNA mutations. There is both evidence of germline selection against and for detrimental mtDNA variants, which raises questions about the potential for positive selection for beneficial mutations within the germline of individuals. Such a mechanism could facilitate mtDNA adaptation, in addition to selection among individuals, especially when the copy-number of mtDNA molecules carrying beneficial mutations within the germline is small. Lastly, given the inherent epistasis required between coadapting mitochondrial- and nuclear-encoded gene products for the maintenance of mitochondrial function, mitonuclear coevolution through adaptive mtDNA changes can contribute to reproductive isolation through the evolution of cytoplasmic incompatibility.

Taken together, our results demonstrate that mitochondrial genomes are not merely passive recipients of nuclear compensation but can themselves be active participants in mitonuclear adaptive evolution. By directly linking mtDNA sequence changes to fitness recovery, ETC-complex specificity, and repeatable genic targets, this study provides experimental validation of a more reciprocal model of mitonuclear coadaptation. These findings have broad implications for understanding evolutionary rescue, the maintenance of mitochondrial function despite high mutation rates, and the molecular bases of mitonuclear incompatibilities relevant to aging, disease, and hybrid dysfunction. More generally, they highlight the power of prospective experimental evolution to disentangle cause and effect in complex coevolutionary systems.

## Methods

### ETC-deficient mutants

Four ETC-deficient mutant alleles of *cox-1*, *ctb-1*, *gas-1* and *isp-1* affecting ETC complexes comprising both mtDNA- and nucDNA-encoded subunits were selected for this study (**Table 1**). In addition to the four single mutants listed above, we also included a double mutant of *isp-1*/*ctb-1*.

The nuclear *gas-1* gene in *C. elegans*, an ortholog of NDUFS2 in humans, encodes a core subunit of ETC complex I where it functions to promote the assembly of the quinone-binding pocket (Vasta et al. 2011; Shiraishi et al. 2012). The *gas-1*(*fc21*) mutant allele was generated by ethyl methanesulfonate (EMS) mutagenesis and is characterized by a nonsynonymous base substitution that replaces a highly conserved arginine with a lysine at amino acid residue 290 within a core subunit of NADH:ubiquinone-oxidoreductase (ETC complex I) (Kayser et al. 1999). This mutant has strongly reduced complex I-dependent metabolism (Kayser et al. 2004), and accumulates oxidative damage in mitochondrial proteins owing to its increased ROS production (Kayser et al. 2001, 2004). The nuclear *gas-1*(*fc21*) mutant allele was obtained from the Caenorhabditis Genetics Center (University of Minnesota) and backcrossed to Bristol N2 for 10 generations to generate an isogenic mutant strain (Wernick et al. 2019).

Mutants of the nuclear-encoded ETC gene *isp-1* exhibit dramatically different phenotypes than those of *gas-1* (**Table 1**; Feng et al. 2001). The gene encodes ISP-1, the Rieske iron sulphur protein of ETC complex III, which transfers electrons from ubiquinol to cytochrome c, and, like ETC complex I, simultaneously pumps protons across the mitochondrial inner membrane thereby helping to establish the proton gradient. ISP-1 and its position within complex III is well suited for studying the evolution of within- and between-genome interactions. This complex consists of several nucDNA-encoded subunits and the mtDNA-encoded cytochrome b (*ctb-1*). *isp-1* mutants exhibit retarded metabolism and extended lifespan (Rea 2005; Ventura et al. 2006). We utilized the *isp-1*(*qm150*) mutant allele bearing a proline to serine replacement in the head domain of ISP-1 (Feng et al. 2001). This region functions to transfer reducing equivalents within complex III to cytochrome c1 through a series of conformational changes (Iwata 1998). Homology modelling indicates that this mutation distorts the structure and alters the redox potential of ISP-1 (Feng et al. 2001, Jafari et al. 2015). Thus, the “slow-living” phenotype of *isp-1* appears to result from an increased reliance upon alternate (and less efficient) metabolic pathways for energy production and a concomitant reduction in mitochondrial respiration (Rea 2005; Jafari et al. 2015). The isp-1(*qm150*) allele has been described as “a healthy mutant with very slow physiological rates” (Feng et al. 2001). However, its delayed maturation and reduced fecundity (98) reduce total and competitive fitness (Dietz et al. 2025). Additionally, *isp-1*(*qm150*) mitochondrial ROS production is elevated (Yang et al. 2007; Lee et al. 2010; Yang and Hekimi 2010) while net ROS levels *in vivo* are not substantially different from wildtype (Dingley et al. 2009) —a difference that appears to be due to increased rates of ROS production and ROS scavenging (Dues et al. 2017). Congruent with this idea, expression of antioxidant and a variety of other stress response genes is upregulated in this mutant (Cristina et al. 2009; Yang and Hekimi 2010; Dues et al. 2017)

The same mutant screen (Feng et al. 2001) that identified the *isp-1*(*qm150*) also yielded an *isp-1*(*qm150*);*ctb-1*(*189*) double mutant which was additionally subjected to the adaptive regime in this study. The Estes laboratory subsequently isolated *ctb-1*(*189*)-bearing mitochondria onto a wildtype N2 nuclear background. *ctb-1*(*qm189*) is a homoplasmic allele that substitutes a valine for a conserved alanine in CTB-1 near the binding site of the ISP head domain (Feng et al. 2001). The *isp-1* and *ctb-1* mutant locations are not predicted to directly interact (Iwata 1998); however, *ctb-1*(*qm189*) partially suppresses the *isp-1*(*qm150*) phenotype via beneficial allosteric effects on complex I (Suthammarak et al. 2009). This finding makes sense in light of the fact that ETC complexes I, III and IV form stable supercomplexes that improve ETC functionality (Acín-Pérez et al. 2008). *isp-1*(*qm150*) weakens the association of this supercomplex and reduces the amount and activity of complex I. The *ctb-1*(*qm189*) mutation exhibits sign epistasis (Weinreich et al. 2005) since it is beneficial within the context of *isp-1*(*qm150*), but in isolation causes slightly deleterious effects on fitness and complex III activity (Suthammarak et al. 2009).

The mtDNA gene *cox-1* encodes cytochrome c oxidase I (COX-1), the main catalytic subunit of cytochrome c oxidase or ETC complex IV. The three mtDNA-encoded subunits (COX-1, 2 and 3) form the functional core of the complex whereas the nDNA-encoded subunits are essential for complex assembly and function (Barrientos et al. 2002; Li et al. 2006). Many additional nucDNA genes are essential for biogenesis of the functional complex (Barrientos et al. 2002), and are thus potential targets of compensatory mutation. ETC complex IV transfers electrons from cytochrome c to reduce oxygen to water. It is not believed to be a major contributor to ROS production; however, its structural/functional state may indirectly affect ROS generation via other members of the I:III:IV supercomplex (Greggio et al. 2017). Cytochrome c oxidase deficiency is a leading cause of human mitochondrial disorders (Shoubridge 2001), and inter-population (Rawson & Burton 2002) and interspecies (Sackton et al. 2003) hybrid incompatibilities are documented to result from breakup of coadapted gene complexes involving this enzyme. The mitochondrial *cox-1* mutation (m.7878G > T) isolated from the CB4856 Hawaiian natural isolate of *C. elegans* strain (Dingley et al. 2014) replaces an alanine with a serine in the N-terminus of COX-1 within the matrix side of the complex IV catalytic core relative to the laboratory N2 strain. This natural variant was purported to be advantageous to CB4856 worms cultured at their native temperature of 25°C, but deleterious for fitness at the standard laboratory temperature of 20°C (Dingley et al. 2014). A transmitochondrial cybrid strain containing the CB4856 mtDNA genome bearing this homoplasmic variant on a N2 nucDNA background was kindly provided by Dr. Marni Falk (Children’s Hospital of Philadelphia).

### Construction of ETC mutant and control strains across three breeding system backgrounds

*C. elegans* is unique among metazoan experimental systems in that sex determination and breeding system can be genetically manipulated to yield populations with variable ratios of hermaphrodites, females and males, thereby achieving different degrees of selfing and outcrossing (Teotónio et al. 2017). In order to investigate the impact of sexual/breeding system on the rates and patterns of mitonuclear adaptation, each of the four single mutant alleles (**Table 1**) initially present on the N2 wildtype (facultatively outcrossing) was isolated onto two additional genetic backgrounds, namely *xol-1* (obligate selfing) and *fog-2* (obligate outcrossing). XOL-1 is a metabolic kinase that functions in X chromosome dosage compensation. The disruption of this function in *xol-1* mutants causes male (X0) lethality, resulting in obligate selfing (Anderson et al. 2010). FOG-2, an F-box protein, helps to initiate sperm production in hermaphrodites (Schedl & Kimble 1988). *fog-2* mutant populations comprise feminized hermaphrodites and males that reproduce via obligate outcrossing. Importantly for this study, mutant *fog-2* populations bearing a loss-of-function allelic substitution that were subjected to laboratory evolution reverted from obligate outcrossing to selfing at a low rate (Katju et al. 2008) via ectopic gene conversion involving a neighboring paralog*, ftr-1* (Rane et al. 2010). Hence, to preclude the conversion of our obligately outcrossing (*fog-2* mutant) lines to selfing during the adaptation regime, we first generated a *fog-2* deletion allele via CRISPR-Cas9 editing (NemaMetrix/InVivo Biosystems, Eugene, OR) to serve as a control strain and for the subsequent creation of obligately outcrossing populations of the ETC-deficient mutants. Similarly, the control *xol-1* mutant strain was generated by CRISPR-Cas9 mediated deletion of the *xol-1* gene in our laboratory stock of wildtype N2. The N2, *fog-2* deletion and *xol-1* deletion strains serve as controls for the effects of the ETC-deficient mutants.

As mentioned above, the four ETC-deficient mutant strains were already available in a wildtype N2 (facultatively outcrossing) genetic background. To generate obligately outcrossing (*fog-2* mutant) populations bearing the four focal ETC-deficient mutations, we established replicate genetic crosses between a hermaphrodite of the N2 strain bearing the ETC-deficient mutation with three males of the control *fog-2* deletion strain to promote outcrossing. Plates with equal male/hermaphrodite sex ratios were selected as the equal sex ratios confirmed the outcrossing of the hermaphrodite with the *fog-2* mutant male. Because *fog-2* mutations are recessive, the F1 offspring comprise hermaphrodites and males. In the case of the homoplasmic mtDNA-encoded ETC mutants (*cox-1* and *ctb-1*), all F1 outcrossed offspring were expected to have inherited the mtDNA mutant allele from the hermaphrodite mother but were heterozygous at the *fog-2* locus (one wildtype and one deletion allele).

Hermaphrodites/females were mated with a male sibling for several additional generations to permit a homozygous mutant genotype at the *fog-2* locus which renders females instead of hermaphrodites. For the nucDNA-encoded ETC mutants (*gas-1* and *isp-1*), all F1 hermaphrodite and male offspring are expected to be double heterozygotes. We repeatedly crossed of hermaphrodites/females with a male sibling for several additional generations until both parents were verified to be homozygous for the *fog-2* deletion and ETC-deficient mutant allele via molecular screening (PCR, electrophoresis and Sanger sequencing).

Because the *xol-1* deletion strain is obligately selfing, additional mutants could not be introduced into this genetic background via standard crossing with another mutant strain. To generate the ETC-deficient mutants in a *xol-1* mutant background, we used CRISPR-Cas9 editing to engineer a *xol-1* deletion in the wildtype N2 strains carrying the ETC-deficient mutation. Repeated selfing of hermaphrodites for several additional generations yielded a strain bearing the focal ETC-deficient mutant and homozygous for the *xol-1* deletion allele, and subsequently verified by PCR and Sanger sequencing. This strategy was successful for generating the *cox-1*, *ctb-1* and *gas-1* ETC mutants in a *xol-1* mutant background. However, due to a failure of introducing the *xol-1* deletion into the *isp-1* mutant line, an *isp-1* mutation was introduced into the *xol-1* deletion strain instead.

The resultant strains served as ancestors of the adaptive RC lines and were cryogenically preserved at −80°C.

### Laboratory adaptation at large population sizes

A total of 16 ancestral strains served as the progenitors for this experimental evolution study. This set comprised of four single ETC-deficient mutants across three breeding systems (4 × 3 = 12 treatments), the double mutant in the N2 wildtype background (1 treatment) and three breeding system control strains lacking an ETC-deficient mutant (N2 wildtype, *xol-1* deletion mutant and *fog-2* deletion mutant = 3 treatments). Eight replicates of each of the 16 ETC ancestral mutant and control strains were experimentally evolved in parallel at large population sizes (i.e., bottleneck sizes of 1,000) for 60 consecutive generations, thereby yielding 128 “recovery lines” (RC lines; cf., Estes and Lynch 2003). The eight *gas-1* mutant lines in the wildtype N2 background were generated in a preceding experiment (Wernick et al. 2019). The adaptive experimental evolution regimes followed previous methods (Wernick et al. 2019; Bever et al. 2022) wherein populations were maintained at 20°C on 100 mm Petri dishes containing Nematode Growth Medium Light (NGML), 1 ml of 200 mg/ml streptomycin, and a bacterial lawn of streptomycin-resistant OP50-1 *Escherichia coli* as a food source. A standardized bleach treatment was applied to each of the 128 RC lines each generation to maintain evolving populations in non-overlapping generations. Each generation, worms were rinsed from crowded plates using M9 buffer and dispensed into 15 ml conical tubes followed by centrifugation at 800 rpm for 30s. The excess M9 supernatant was discarded while leaving the worm pellet intact at the tube bottom. A mixture of three parts diluted commercial bleach (final concentration = 2.75% bleach in ddH2O) and one part 5M NaOH was then added to the conical tubes. Tubes were inverted every 2min to dissolve the worms while releasing viable embryos followed by centrifugation to form an embryo pellet. The bleach solution was discarded and followed by three rounds of rinsing with fresh M9 buffer. The embryo pellet was then transferred to a 1 ml microtube and vortexed, after which 1 ul was transferred onto an eight-well slide. Embryo counts were used to calculate the amount needed to transfer 1,000 individuals onto new large plates to initiate the next generation. Strains were transferred when the majority of hermaphrodites or females reached peak gravidity and began laying embryos with a few hatched larvae; plates were typically well-starved by this point. Following 60 generations of maintenance at expanded population sizes, stocks of the adaptive RC lines were cryogenically preserved at −80°C.

### Fitness assays

The fitness assays were conducted following standard protocols as previously reported (Dietz et al. 2025). We assayed fitness-related traits for all lines alongside the wildtype N2 control and the appropriate ancestral mutant. For all facultatively outcrossing (N2 background) and obligately selfing (*xol-1* background) lines, we assayed daily production of selfed progeny following established methods (e.g., Wernick et al. 2019). These assays were initiated by allowing 10–15 adult hermaphrodites from a line to lay embryos for 5h. Single embryos were then transferred to individual 60 mm Petri plates containing NGML, 1 ml of streptomycin, and OP50-1 *E. coli* food source, and allowed to develop. Once hatched, the number of plates was reduced to 20 for the N2 control, 10 for each ancestral mutant, and five for all RC lines following 60 generations of population expansion. At the same time each day, hermaphrodite parents were transferred to a fresh plate. Offspring were allowed to develop to the L3/L4 larval stage and then killed with a drop of 0.5 M sodium azide and stored at −4 °C to be counted. Offspring were counted by counterstaining plates with toluidine blue dye.

Outcrossed progeny production was assayed for the obligately outcrossing *fog-2* lines. These assays were initiated by picking individual L4 larval stage male and female pairs onto fresh plates–20 pairs for the ancestral mutant and 10 pairs for all RC lines following 60 generations of population expansion. The focal pairs were transferred together every 24h and offspring were counted as described above.

Offspring counts from both selfed- and outcrossed-fitness assays were used to generate reproductive schedules and calculate total reproductive output and relative fitness of the mutants compared with N2, and with the relevant ancestral control following Christy et al. (2017). Relative fitness of each individual was computed as: *ω* = *Σe ^-rx^ l(x)m(x)*, where *l(x)* is the number of worms surviving to day *x*, *m(x)* is the productivity (number of surviving progeny) at day *x*, and *r* is the mean intrinsic population growth rate of the assay-specific N2 or ancestral mutant control as appropriate. The latter was calculated by solving Euler’s equation for *r* from *ω* = *Σe ^-rx^ l(x) m(x)* = 1 using an average value of *l(x) m(x)* for each block-specific control. We used *x* = 4.75 on the first reproductive day (cf., Vassilieva et al. 2000).

### Genomic DNA extraction and whole genome sequencing of adaptive recovery lines

Cryopreserved stocks of the 128 adaptive RC lines following 60 generations of experimental evolution were thawed to establish large starting populations (>200 worms) for tissue collection. Thawed populations were established on standard 90 mm NGM plates seeded with a lawn of *E. coli* OP50 and incubated at 20°C. For each line, nematode populations were allowed to grow for an additional two to three generations and then collected with M9 solution and transferred to two large 90 mm NGM plates seeded with *E. coli* OP50 to enable further population expansion. Worm tissue was collected in 0.5 ml TEN solution and frozen at -20°C for subsequent genomic DNA extraction. Genomic DNA for each line was isolated from the frozen tissue using the PureGene Genomic DNA Tissue Kit (Qiagen no. 158622) as per a previously described protocol (Katju et al. 2022). DNA quality and quantification were assessed via electrophoresis on 1% agarose gels, a Thermo Fisher Nanodrop spectrophotometer and BR Qubit assay (Invitrogen). For each adaptive RC line, 2 ug of high-quality gDNA was used to generate genomic libraries yielding 150 bp paired-end reads. Whole genome sequencing was conducted on multiplexed libraries by 2×150 paired- end runs using Illumina NovaSeq 6000 at the AgriLife Genomics & Bioinformatics Center (Texas A&M University) and aiming for >50× and >1,000× coverage for the nDNA and mtDNA genomes, respectively.

### Preprocessing and mapping of reads

The quality of the raw sequencing reads was assessed using the software FastQC v0.11.9 (Andrews 2010). Adapter trimming and low-quality base filtering were performed using Trimmomatic v0.39 (Bolger et al. 2014), 5′ and 3′ low-quality trailing bases and reads shorter than 75 bp were removed. The filtered reads were then aligned to the *C. elegans reference* genome (N2 strain, NCBI reference WBcel235, WormBase release WS295) using BWA-MEM v0.7.18 (Li & Durbin 2009) with default parameters. The resulting files were converted to BAM format, sorted, indexed and a minimum mapping quality filter of 10 was applied using the samtools suite v1.20 (Li et al. 2009). Finally, PCR and optical duplicates were tagged using the Picard Tools v3.1.1 *MarkDuplicates* function.

### Variant calling, quality control and annotation

Small variant calling was performed in parallel with BCFtools mpileup (Li 2011) and GATK v.4 following the suite’s best practices (McKenna et al. 2010). Copy-number variants were assessed based on depth variation and read mapping abnormalities through the use of CNVnator v0.4.1 (Abyzov et al. 2011) and Lumpy (Layer et al. 2014). All background-specific mutations were removed to retain the unique mutations. Due to the low number of detected mutations, variant quality control was conducted manually by inspecting alignments in IGV v2.18.4 (Robinson et al. 2011). Variants with coverage as low as 2% were retained to ascertain low-frequency heteroplasmic mutations. The final high-confidence variants were annotated using SnpEff v5.2c (Cingolani et al. 2012), using the WormBase reference database WBcel235.

### 3D protein structural locations of high-frequency mtDNA variants

AlphaFold3 (Abramson et al. 2024) was used to map the protein structural locations of 11 mutated amino acid residues resulting from putative compensatory high-frequency nonsynonymous mutations in a subset of *gas-1* and *isp-1* adaptive RC lines is relation to the focal deleterious mutation. These included (i) eight nonsynonymous changes in *gas-1* (ETC complex I) adaptive RC lines (four each in *nd-1* and *nd-6*), and (ii) three nonsynonymous changes localizing to *ctb-1* in *isp-1* (ETC complex III) adaptive RC lines.

### Alternative open reading frame (altORF) identification and analysis

We followed the general methods of Kienzle et al. (2023) to interrogate the alternative mitochondrial proteome of the wildtype *C. elegans* mtDNA reference sequence, GenBank accession number NC_001328.1. First, we used a custom Python script and the ORF finder tool within Geneious Prime Version 2023.2.1 (https://www.geneious.com) to conduct an *in silico* search for ORFs across all six reading frames, specifically focusing on those encoding peptides of 20 amino acids or longer. The search utilized NCBI’s translation table 5, considering alternative initiation codons relevant to *C. elegans*. We excluded interior ORFs and end codon positions associated with known protein-coding genes. Second, we used the OpenProt database (Leblanc et al. 2024), which relaxes traditional annotation criteria and may offer a more biologically relevant perspective of the mitochondrial proteome. Finally, we mapped and compared the locations of the identified altORFs to those of the mtDNA variants identified in our sequenced recovery lines using Geneious Prime.

## Data and Code Availability

Whole mtDNA genome sequence data will be deposited at NCBI and will be made available upon acceptance. All code will be made available online at Github.com.

## Acknowledgements

This work was funded by National Science Foundation awards MCB-2232413 to S.E., MCB- 1817762 to V.K. and U.B., and MCB- 1817993 to V.K., U.B. and S.E., as well as an American Heart Association predoctoral fellowship (23PRE899291) and Portland State University Forbes-Lea research grant to Z.P.D. J.L.C. was sponsored by a fellowship from the Carl Trygger Foundation grant to V.K. The computations were enabled by resources provided by the National Academic Infrastructure for Supercomputing in Sweden (NAISS) and the Swedish National Infrastructure for Computing (SNIC) at Uppmax [partially funded by the Swedish Research Council through grant agreements no. 2022-06725 and no. 2018-05973].

## Author Contributions

V.K., U.B. and S.E. conceived the original experimental idea. V.K. constructed the ancestral strains. S.E. evolved 128 adaptive recovery lines for 60 generations. V.K. and U.B. oversaw gDNA collection from the lines and submitted samples for whole-genome sequencing. V.K. and U.B. designed the analyses of the data. V.K. carried out most analyses with some supplementary analyses performed by U.B., J.L.C. and Z.P.D. The manuscript and figures were drafted by V.K. and U.B. with some supplementary sections written by J.L.C. and Z.P.D. and improved with help from S.E. All authors revised and approved the article.

