## Supplementary Material for "No passive sidekick: mitochondrial compensatory evolution during adaptation of ETC mutant lines in *Caenorhabditis elegans*"

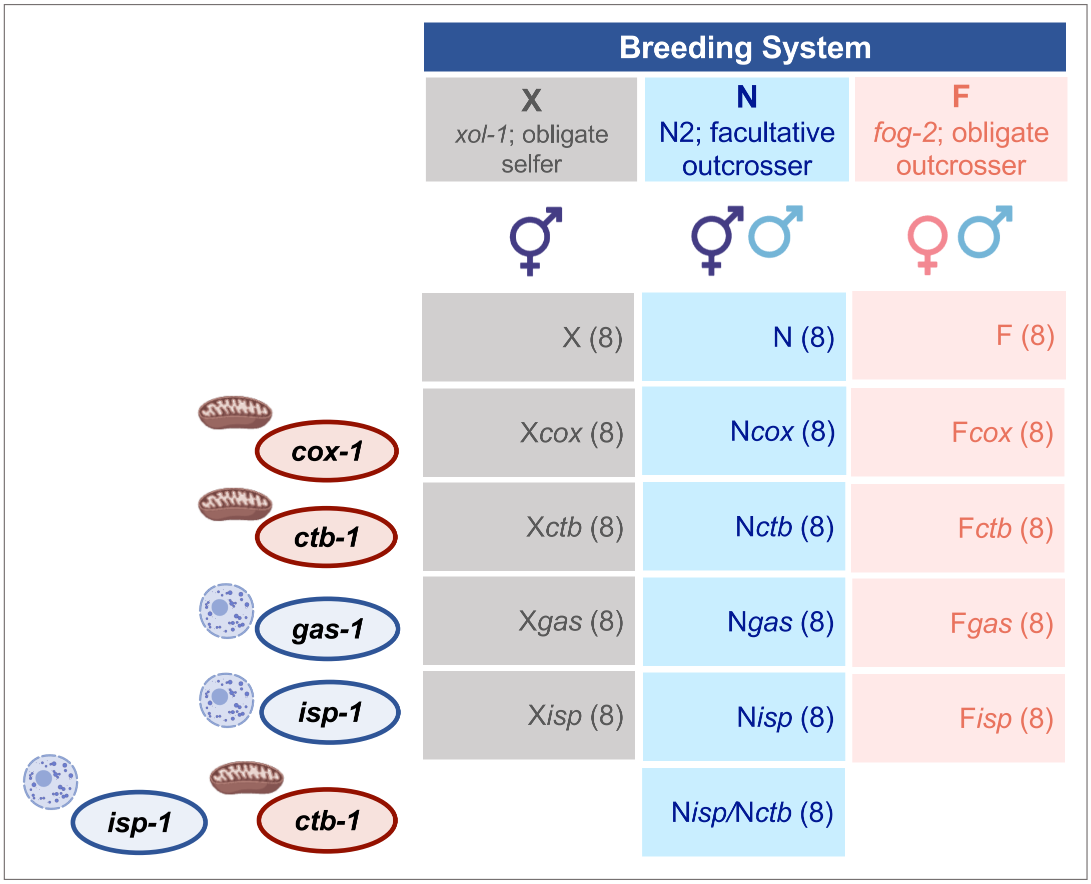


**Supplementary Fig. S1. Experimental design and line notation**. Four electron transport chain (ETC) mutants, two mtDNA-encoded (*cox-1* and *ctb-1*) and two nuclear-encoded (*gas-1* and *isp-1*) introduced into three different breeding systems. The three breeding systems denoted N, F and X represent variable ratios of males, females, and hermaphrodites, and therefore varying degrees of selfing and outcrossing to allow direct tests of the role of recombination in evolution. The four ETC mutants in the three breeding system backgrounds and an additional facultatively outcrossing double mutant (*isp-1*/*ctb-1*) represent 13 treatments. The three breeding systems without an ETC mutant served as large population size controls. and comprise an additional three treatments. Eight biological replicate lines were evolved for each of the 16 treatments, totaling 128 experimental evolution lines.

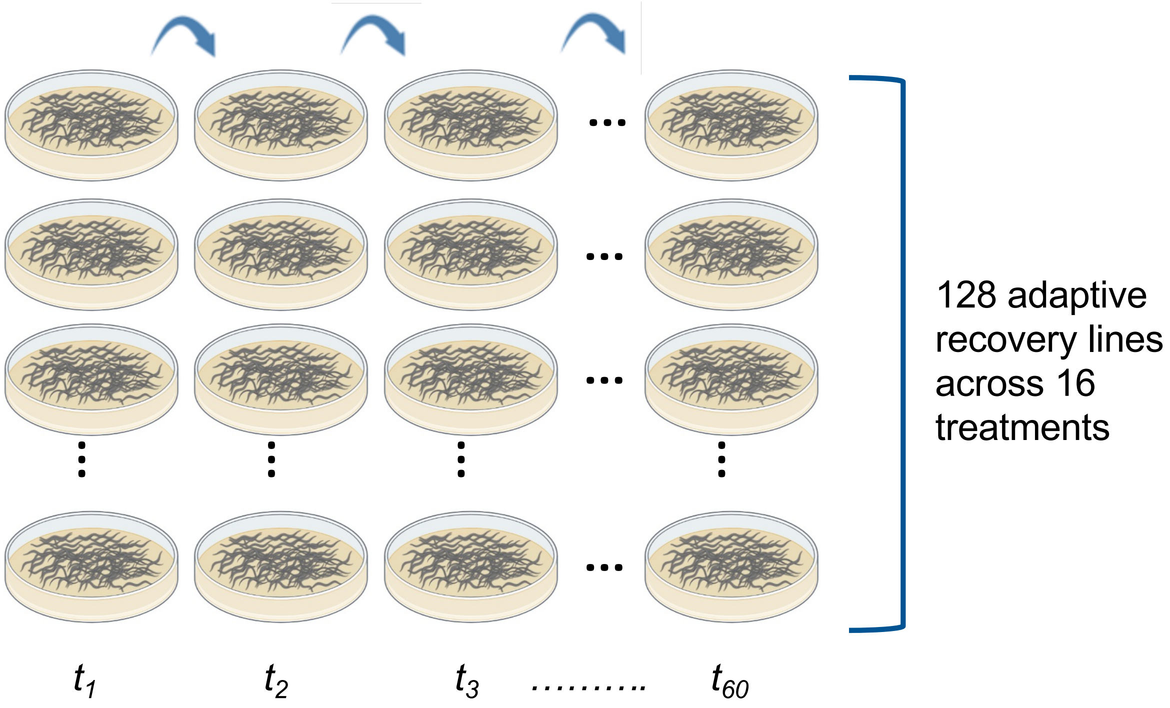


**Supplementary Fig. S2. Adaptive recovery (RC) experiment.** To enable fitness/adaptive recovery of the ETC mutant and control lines, the 128 experimental lines were expanded and independently maintained at large population sizes for 60 consecutive generations.

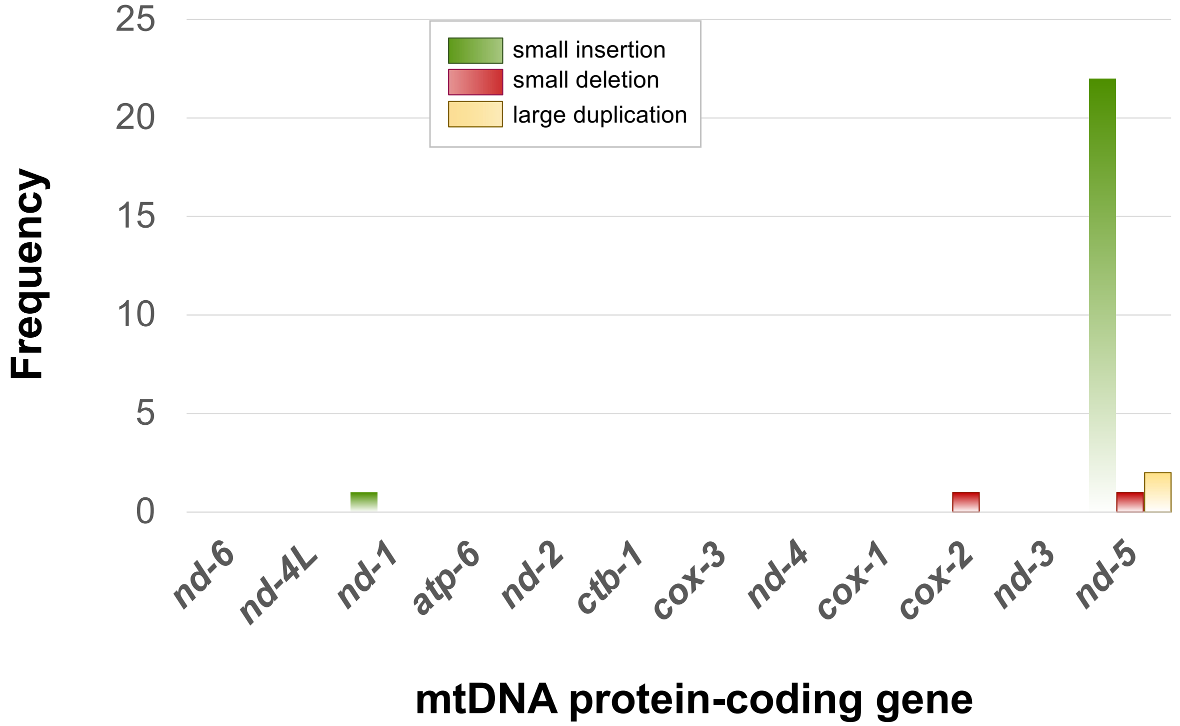


**Supplementary Fig. S3. Indels per protein-coding gene**. Frequency distribution of novel insertion and deletion mutations in the adaptive RC lines across the 12 protein-coding genes comprising the *C. elegans* mtDNA genome. The indels are further classified into small insertions (≤ 3 bp; red), small deletions (≤ 3 bp; green), and large duplications (yellow) and deletions. No large deletion variants were identified in the lines following 60 generations of adaptive RC.

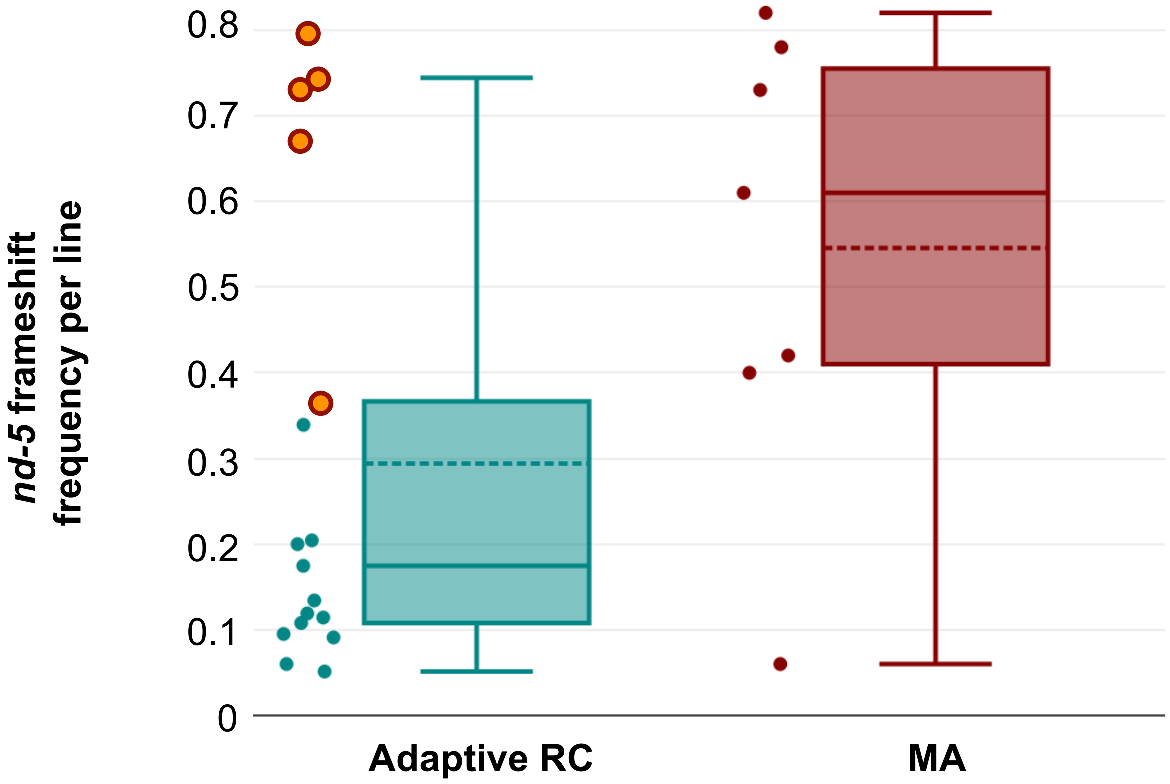


**Supplementary Fig. S4. Frequencies of *nd-5* frameshift mutations in adaptive RC lines *vs.* MA lines**. The majority of *nd-5* frameshift mutations were in <20% frequency in the RC lines (left) except for the double mutant *isp-1*/*ctb-1* RC lines (orange circles) where their frequencies were more similar to the high-frequency *nd-5* frameshift mutations found in MA lines (right) (Konrad et al. 2017).

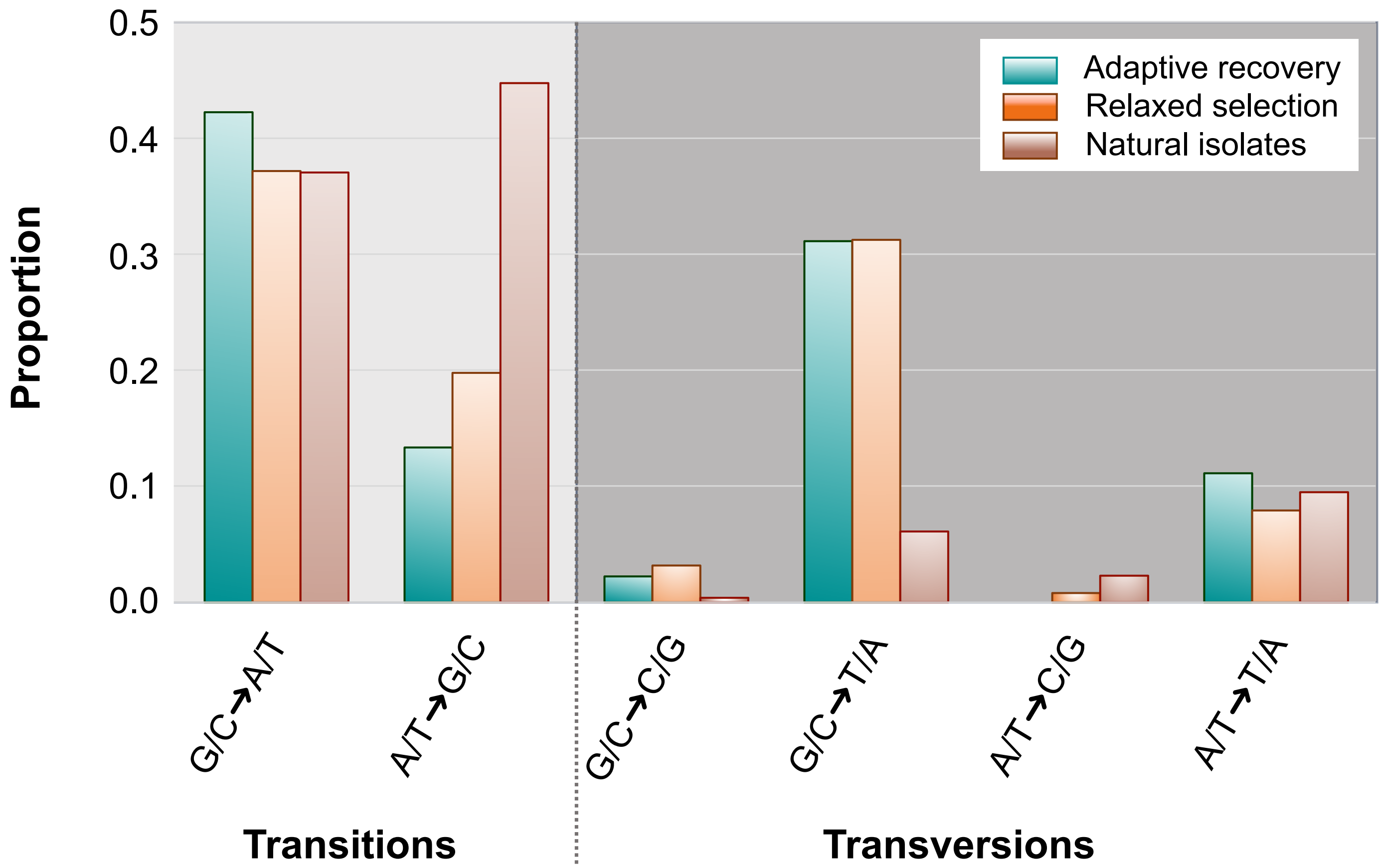


**Supplemental Fig. S5. The mutational spectrum of base substitutions in the mitochondrial genomes of *C. elegans* adaptive RC lines.** The proportion of base substitutions are normalized by the frequencies of each individual variant. The spectrum detected in our adaptive RC lines is contrasted with the spectrum of mtDNA base substitutions observed in preceding studies of *C. elegans* MA lines (Konrad et al. 2017) and *C. elegans* natural isolates (Schifano et al. 2026).

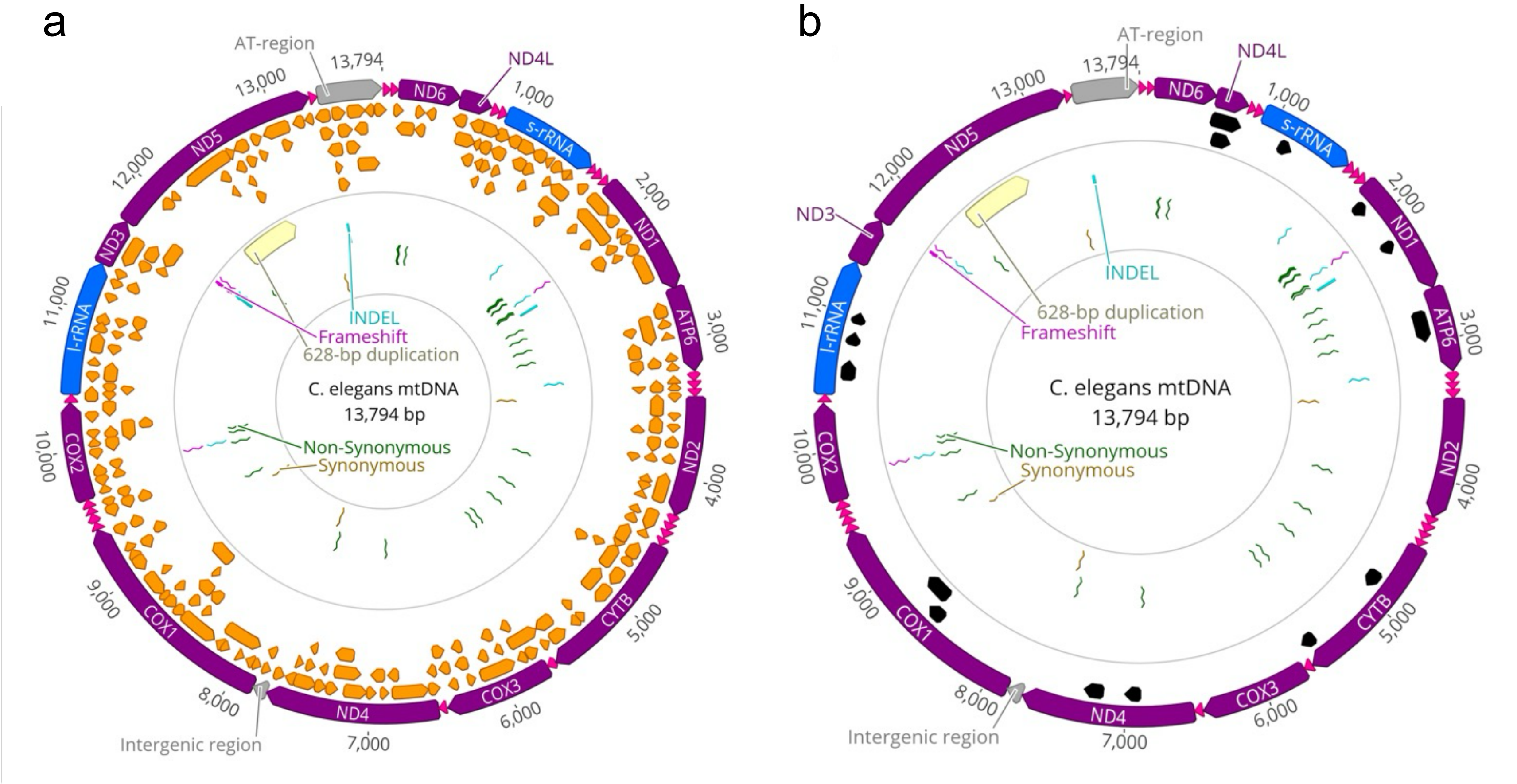


**Supplementary Fig. S6. Potential altORFS and smORFs in the *C. elegans* mitochondrial genome. a,** Potential small ORFs and alternative ORFs in the *C. elegans* mitochondrial genome as identified in silico. In purple: the typical 12 protein-coding genes. In blue: the l-rRNA and s-rRNA genes. In pink: the various tRNA genes. In orange: Arrows positioned inside the circle indicate smORFs and altORFs on the main coding and complementary strands in all six reading frames. In green, gold, light blue, magenta, and yellow: the location of nonsynonymous, synonymous, small indels, frameshift mutations and a 628 bp-duplication, respectively, as identified in **Supplementary File 1**. **b,** Potential alternative ORFs in the *C. elegans* mitochondrial genome as identified with OpenProt. In purple: the typical 12 protein-coding genes. In blue: the l-rRNA and s-rRNA genes. In pink: the various tRNA genes. In black: Arrows positioned inside the circle indicate 10 altORFs on the main coding and complementary strands in all six reading frames as identified with OpenProt. In green, gold, light blue, magenta, and yellow: the location of nonsynonymous, synonymous, indel mutations, frameshift mutations and a 628 bp-duplication, respectively, as identified in **Supplementary File 1**.
